# The basolateral amygdala dopamine system shapes dynamic drug seeking states

**DOI:** 10.64898/2026.09.29.755395

**Authors:** Megan A. Brickner, William E. Szot, Julianna M. Prohofsky, Amy R. Wolff, Mark J. Thomas, Benjamin T. Saunders

## Abstract

Drug-associated sensory cues and contexts exert powerful influence over drug seeking motivation, but in vivo brain mechanisms governing this process remain unclear. Here, we recorded and manipulated dopamine signaling and dopamine receptor D1+ neuronal activity in the basolateral amygdala (BLA) to investigate their contributions to dynamic sensory-guided cocaine seeking in rats. First, we show that cocaine exposure sensitizes dopamine transmission from ventral tegmental area dopamine neurons projecting to the BLA. We then employed an intermittent access cocaine self-administration paradigm that prompted robust binge-like cocaine seeking that was heavily guided by shifts in the sensory environment. Cued transitions to drug available states evoked large and sustained BLA dopamine signals and transient neuronal responses from BLA D1+ neurons. BLA dopamine signals and D1-neuron activity came to preferentially encode these drug availability states, which hierarchically modulated neural activity during drug seeking actions and predicted different aspects of behavior. Finally, we found that optogenetically enhancing BLA D1-neuron activity at drug availability state transitions potentiated cocaine binging. Our findings provide new insight into the neural control of addiction-related behavior, identifying the dopaminergic amygdala as a key interface between the environment and decisions to engage in vigorous drug use.

## INTRODUCTION

Craving, drug seeking, and relapse in addiction are heavily controlled by drug-associated sensory “cues” (sights, sounds, places) that dynamically spur motivation and reinforce actions (Robinson and Berridge, 1993; Everitt and Robbins, 2005; Perry et al., 2014; Namba et al., 2018; LeCocq et al., 2020; Lüscher and Janak, 2021). This sensory landscape is complex, including global or contextual cues that signal states of drug availability, such as seeing a “smoking area” sign, and also proximal cues associated with drug delivery, such as holding a cigarette (Weiss et al., 2000; Chaudhri et al., 2008; Crombag et al., 2008). Environmental cues often signal intermittent access to drugs, which promotes key addiction-related behaviors, including binge drug use (Crombag et al., 2008; Zimmer et al., 2012; Calipari et al., 2013, 2015; Kawa et al., 2016, 2019; Allain and Samaha, 2019; Fraser and Holland, 2019; Minogianis et al., 2019; Collins et al., 2023; Mainville-Berthiaume et al., 2026).

Despite the strong theoretical background and translational implications, we still know relatively little about in vivo neural mechanisms that govern sensory-guided drug seeking motivation. While much existing literature has focused on striatal dopamine functions (Everitt and Robbins, 2005; Aarts et al., 2011; Yager et al., 2015; Poisson et al., 2021), the basolateral amygdala (BLA) is positioned to integrate sensory and emotional inputs to guide cue learning and dynamic affective states in addiction (Wassum and Izquierdo, 2015). In addition to a variety of glutamatergic inputs, the BLA receives robust dopamine neuron input from the ventral tegmental area (VTA) (Haber and Fudge, 1997; Brinley-Reed and McDonald, 1999; Takahashi et al., 2010; Mingote et al., 2015) and there is evidence implicating the BLA dopamine system in drug seeking (Wilson et al., 1994; Hurd et al., 1997; See et al., 2001; Carelli et al., 2003; Di Ciano and Everitt, 2004; Stefanik and Kalivas, 2013; Murray et al., 2015; Kim and Lattal, 2019; Mahler et al., 2019; Puaud et al., 2021; Smith and Torregrossa, 2024). BLA dopamine levels scale in magnitude during valenced events and predictive cues suggesting that conditioned dopamine transmission in the BLA signals the saliency of transitions into a valenced state (Correia and Goosens, 2016; Lutas et al., 2019, 2022; Tang et al., 2020; Grove et al., 2022; Sias et al., 2024; Brickner et al., 2026). Despite growing interest in the circuit, we know little about how the BLA dopamine system is engaged in vivo during the development of addiction-related behaviors.

Here, we interrogated the BLA dopamine system as a locus of drug seeking control. We used a combination of fiber photometry and optogenetics to characterize BLA dopamine and D1-neuron calcium signals as rats engaged in intermittent access cocaine self-administration, which promotes vigorous, sensory-guided drug seeking. Together, our studies reveal a neurobiological mechanism for environmental regulation of drug-seeking motivation, pinpointing the dopaminergic amygdala as a key system for the development of addiction-related behaviors.

## RESULTS

### Cocaine sensitizes VTA to BLA dopamine transmission

We first investigated the impact of cocaine exposure on BLA dopamine transmission evoked by two forms of uncued stimuli: optogenetic activation of VTA dopamine neurons and footshock, both of which engage strong BLA dopamine signaling (Lutas et al., 2019; Tang et al., 2020; Brickner et al., 2026). To do this, we paired in vivo fiber photometry recording of the dopamine dLight1.3b sensor in the BLA with a Cre-driven Chrimson opsin targeted to VTA dopamine neurons in TH-cre rats (**Fig 1A-B**; n=6).

**Fig 1.**
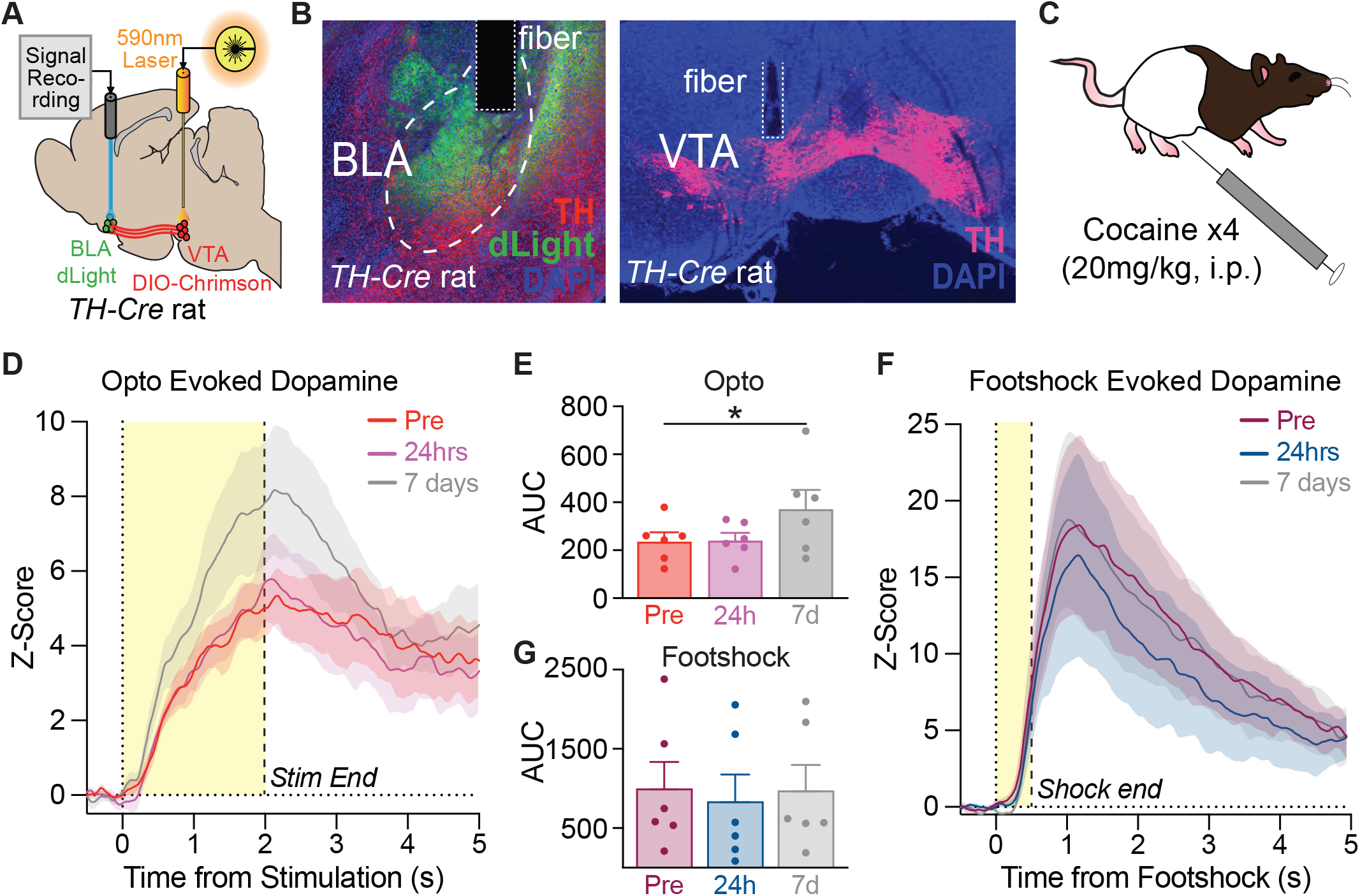
Cocaine sensitizes VTA to BLA dopamine transmission. A) Viral approach for simultaneous in vivo optogenetic activation of VTA dopamine neuron cell bodies paired with fiber photometry recording of dopamine transmission in the BLA via dLight1.3b. Dopamine neurons were targeted in TH-cre transgenic rats with a cre-driven (DIO) Chrimson opsin (n=6, 2F, 4M). B) Histology images showing VTA dopamine terminals over the BLA (left) and VTA dopamine neuron cell bodies (right) expressing TH-mCherry in the VTA. C) Non-contingent cocaine exposure included daily cocaine intraperitoneal injections administered for 4 consecutive days. D) Optogenetic excitation of VTA dopamine cell bodies (2s, 20Hz, 5ms pulses) resulted in robust dopamine release in the BLA before and after cocaine exposure. E) Area under the curve (AUC) measurements of the 2s stimulation window revealed optogenetically evoked VTA to BLA dopamine release was larger 7 days after the final cocaine treatment relative to baseline, pre-cocaine or 24 hours after the final cocaine dose. F) Uncued footshock events triggered large dopamine release into the BLA during each recording session: pre-cocaine, 24 hours after and 7 days after the final cocaine dose. G) AUC measurements of the 2s following footshock onset did not reveal any modulatory effect of cocaine exposure on footshock-evoked dopamine. Data represent mean ± SEM. *p<0.05. i.p.: intraperitoneal.

Optogenetic (590mn laser, 20Hz, 2 sec stim) and footshock (0.4mA) evoked dopamine signals were measured at three timepoints: a pre-cocaine baseline, 24-hours following cocaine sensitization (4 daily intraperitoneal injections, 20mg/kg/injection), and 7-days following cocaine sensitization. Optogenetic activation of VTA dopamine neurons evoked robust dopamine signals in the BLA at the pre-cocaine baseline test, which was similar to the signal 24-hours after cocaine sensitization (**Fig 1D,E**; (p=0.9965). However, 7 days post sensitization, optogenetically evoked dopamine signals were significantly larger (**Fig 1D,E**; repeated measure, one-way ANOVA, F(2,10)=5.729, p=0.0220), relative to both pre-cocaine evoked levels (p=0.0343) and 24-hours post cocaine levels (p=0.0391).

This sensitization effect was specific to optogenetically-evoked dopamine, as we saw no difference in footshock-evoked dopamine in the BLA (**Fig 1F,G**; repeated measure, one-way ANOVA, F(2,10)=0.7641, p=0.4911). This was true for the pre-cocaine versus 24-hours (p=0.5069) and 7-days (p=0.9816) timepoints. These data indicate that cocaine exposure engages VTA-BLA dopamine circuitry to bolster signaling in response to specific inputs, and this effect incubates over time.

### Intermittent access to cocaine promotes binge-like cocaine seeking behavior that is heavily controlled by a shifting environment

We next interrogated how the BLA dopamine system is engaged in vivo during addiction-related behaviors, using a translationally relevant cocaine self administration paradigm that promotes sensory-guided binging cycles of drug use (Calipari et al., 2013; Kawa et al., 2019; Collins et al., 2023). To begin, rats (n=19) learned to self-administer cocaine during continuous access sessions on a FR-1 schedule of reinforcement, where active nose pokes produced cocaine infusions paired with drug delivery cues (**Fig 2A**). After an initial burst of responding on the first session that stabilized thereafter (**Fig 2E**; repeated measures, one-way ANOVA, F(4,72)=9.480, p<0.0001), rats came to discriminate between active and inactive ports by the end of the continuous access phase (**Fig 2F**; two-tailed t-test, t(18)=3.104, p=0.0061). The data in **Fig 2** reflect the dLight and D1-GCaMP cohorts collapsed together, as we saw no differences in cocaine self administration behavior between biosensor groups (**Fig S2**).

**Fig 2.**
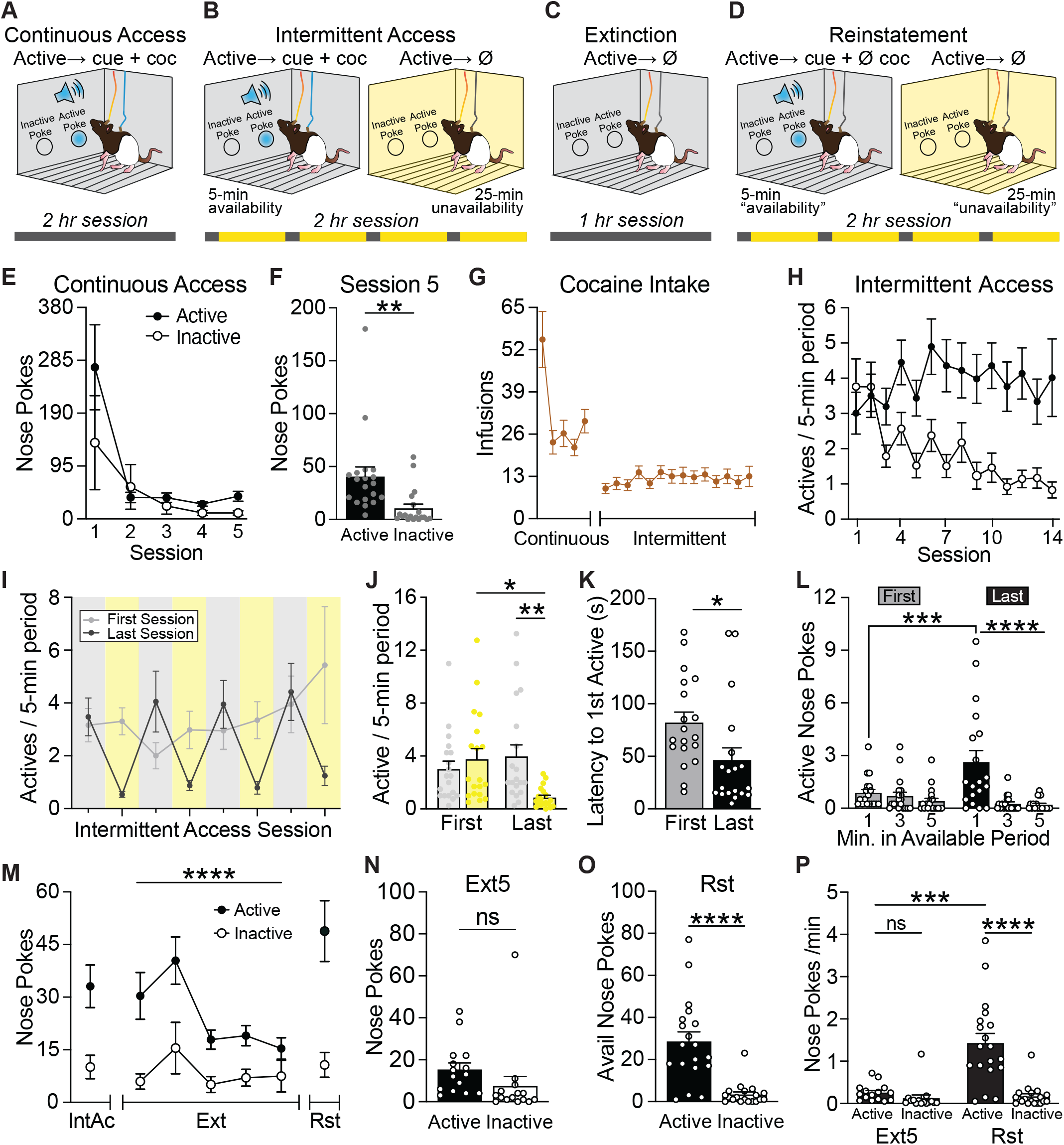
Intermittent access to cocaine prompts dynamic, binge-like drug use bouts controlled by environmental state transitions. (A) Rats (n=19, 10F, 9M) learned to self-administer cocaine with continuous access with the chamber lighting off. Each active nose poke resulted in a drug-delivery cue (nose poke light + tone) and an intravenous cocaine infusion (0.4 mg/kg) followed by a 20s timeout. Rats then underwent sessions of intermittent access to cocaine. 5-min drug available periods were signaled by the chamber lights extinguishing, where active pokes prompted cocaine and cue delivery. The house lights then reilluminated signaling the 25-min drug-free period. There were 4 cycles of drug available and unavailable periods each session. C) Extinction training occurred with the chamber lighting off for the entire session, and nose pokes had no consequences. D) During reinstatement, the cycles of availability states and the drug-delivery cues were reintroduced, but no cocaine was delivered. E) Active (closed circles) and inactive nose pokes (open circles) plotted across continuous access pretraining. F) Late in continuous access, rats make more active nose pokes than inactive. G) Cocaine infusions earned during the continuous access and intermittent access phases. H) Active pokes increased during the drug available periods and decreased in the drug unavailable periods as training progressed. I) Active nose pokes for the first and last Intermittent access to cocaine training sessions (day 1 and day 14) are shown. Rats came to focus the majority of their active nose poke bouts to the available period. Gray boxes represent drug available periods (lights off), while yellow boxes represent drug unavailable (lights on). Gray data points represent early training and black data points represent late training. J) Average active pokes within the 5-min drug available period compared to average active pokes in the unavailable periods binned into 5-min blocks within the first and last intermittent access sessions. K) Average latency to first active poke within the drug available periods decreased between session 1 to session 14. L) Active nose pokes made in minutes 1, 3, and 5 of the drug available periods during the first (gray) and last (black) intermittent access days. M) Comparisons of the active (closed circles) and inactive (open circles) nose pokes sought during the final Intermittent access session, across the 5 extinction sessions, and the conditioned reinstatement test. N) Active and inactive pokes on the final extinction day. O) Comparison of the active and inactive nose pokes made only during the “drug availability” periods during reinstatement. P) Rate of nose pokes per minute of “availability” on the final extinction day and reinstatement. Data represent mean ± SEM. *p<0.05, **p<0.01, ***p<0.001, ****p<0.0001. Ext: extinction. Rst: reinstatement.

Rats then underwent 14 daily sessions of intermittent access training (**Fig 2B**). Within each session, there were 4 cycles of drug available and unavailable periods. Drug available periods were signaled by a global sensory shift, the extinguishing of the chamber lights, which remained off for the entire availability period. Within drug available periods, active nose pokes resulted in an intravenous cocaine delivery and coincident drug-delivery cue. After 5 min of drug availability, the house lights re-illuminated, signaling a 25 min drug-free period where nose-pokes had no consequences. This periodic access to cocaine produced a “saw-tooth” binge-like drug intake pattern where rats rapidly took several cocaine infusions during brief availability (Fig 2G,I). Rats learned to discriminate between availability states, with the majority of nose poking occurring during the drug available periods (**Fig 2H**; main effect of period F(1,18)=42.19, p<0.0001; session x period interaction F(13,216)= 8.520, p<0.0001). This result was further highlighted when specifically comparing the first and last intermittent access sessions (**Fig 2J**; main effect of period F(1,18)=7.926, p=0.0115; session x period interaction F(1,18)=11.31, p=0.0035). Active pokes during the drug unavailable periods diminished between the first and last intermittent access session (p=0.0106) and were significantly lower than active pokes during the drug available period in the final session (p=0.0061).

Across intermittent access training there was a temporal shift in how rats distributed their cocaine intake within availability windows. The latency to first active poke decreased significantly from the first session to the last session (**Fig 2K**; two-tailed t-test, t(18)=2.151, p=0.0453). Intermittent access to cocaine also gave rise to brief, vigorous drug-seeking bouts (**Fig 2L**; main effect of minute F(2,36)=15.21, p<0.0001; minute x session interaction F(2,36)=9.377, p=0.0005), where the majority of active pokes occurred within the first minute of the drug available period (**Fig 2L**; min1 vs min3: p<0.0001; min1 vs min5: p<0.0001), and eventually ceased during the unavailable period. Finally, active pokes during the first minute of the availability period increased from the first and last intermittent access session (p=0.0010) as rats accelerated their drug seeking. Overall, these data indicate that the intermittent transition to a state of cocaine availability promotes vigorous, binge-like drug intake.

### Reintroduction of dynamic availability signals triggers reinstatement of vigorous seeking behavior following extinction

After the intermittent access phase rats underwent extinction, wherein we removed the intermittent transitions signaling drug availability. Each one-hour session began with a single transition into the “drug availability” state signaled by chamber lights turning off for the entire session, but active nose pokes no longer resulted in a cocaine infusion nor the drug delivery cue (**Fig 2C**). Across extinction training, active responses selectively decreased (**Fig 2M**; mixed-effects ANOVA, main effect of session: F(4,56)=8.850, p<0.0001; session x nose poke interaction: F(4,50)=2.628, p=0.0453) with active and inactive nosepokes at similar low levels by the final session (**Fig 2N**; two-tailed t-test, t(14)=1.612, p=0.1294).

After the final extinction session, we conducted a reinstatement test to probe the effect of reintroducing intermittent “drug availability” shifts on reinvigorating drug-seeking behaviors (**Fig 2D**). In this session, 5 min lights off “drug available” periods were intermingled with 25 min lights on periods, mimicking the pattern during intermittent access training. Nose pokes during the lights off period coincided with the drug delivery cues, but no cocaine. This resulted in reinstatement of drug seeking behavior where rats made more active nose pokes than inactive nose pokes during the “availability” periods (**Fig 2O**; two-tailed t-test, t(18)=6.430, p<0.0001). When comparing the rate of nose pokes per minute of “availability” (**Fig 2P**; mixed-effects ANOVA, main effect of phase: F(1,18)=28.84, p<0.0001; phase x nose poke interaction: F(1,10)=24.19, p=0.0006), rate of active (p=0.0001), but not inactive nose pokes (p=0.8696) were significantly higher than on the final day of extinction. This indicates that the reintroduction of dynamic state transitions is sufficient to strongly renew drug seeking motivation that had been previously extinguished in a static environment.

### Sustained BLA dopamine signals encode drug availability state transitions

Using in vivo fiber photometry with a dopamine biosensor (**Fig 3A,B**), we investigated how the BLA dopamine system was engaged during intermittent access to cocaine self administration, focusing initially on signals surrounding the transitions into and out of drug availability states (**Fig 3C**). Dopamine signals in response to either transition were initially small and similar, but as intermittent access training progressed (**Fig 3D-F, Fig 3G-I**), they came to strongly discriminate between drug availability and unavailability transitions based on peak (**Fig 3J**; two-way ANOVA, session x state type interaction: F(2,18)=7.587, p=0.0041; main effect of state type: F(1,9)=11.78, p=0.0075) and area under the curve (AUC) measures of the first 5s after transition (**Fig 3K**; two-way ANOVA, session x state type interaction: F(2,18)=9.753, p=0.0014; main effect of state type: F(1,9)=10.74, p=0.0096), particularly by the middle (peak: p<0.0001; AUC: p<0.0001) and late training phases (peak: p=0.0006; AUC: p=0.0008). As drug seeking became more heavily controlled by the intermittent state transitions, dopamine in response to the availability transition grew larger (**Fig 3D-F**) relative to the early session in middle (peak: p=0.0001; AUC: p<0.0001) and late training (peak value: p=0.0247; AUC: p=0.0391), while the unavailability transition signals stayed low (**Fig 3G-I**). The dopamine responses to the onset of drug availability became sustained, remaining elevated for approximately 20s following the state transition (**Fig S4A-D**). Notably, the average latency to first response (**Fig 2K**) was longer than this 20s window where dopamine is most active, suggesting that this elevation in signal is unlikely to reflect responses to active cocaine consumption.

**Fig 3.**
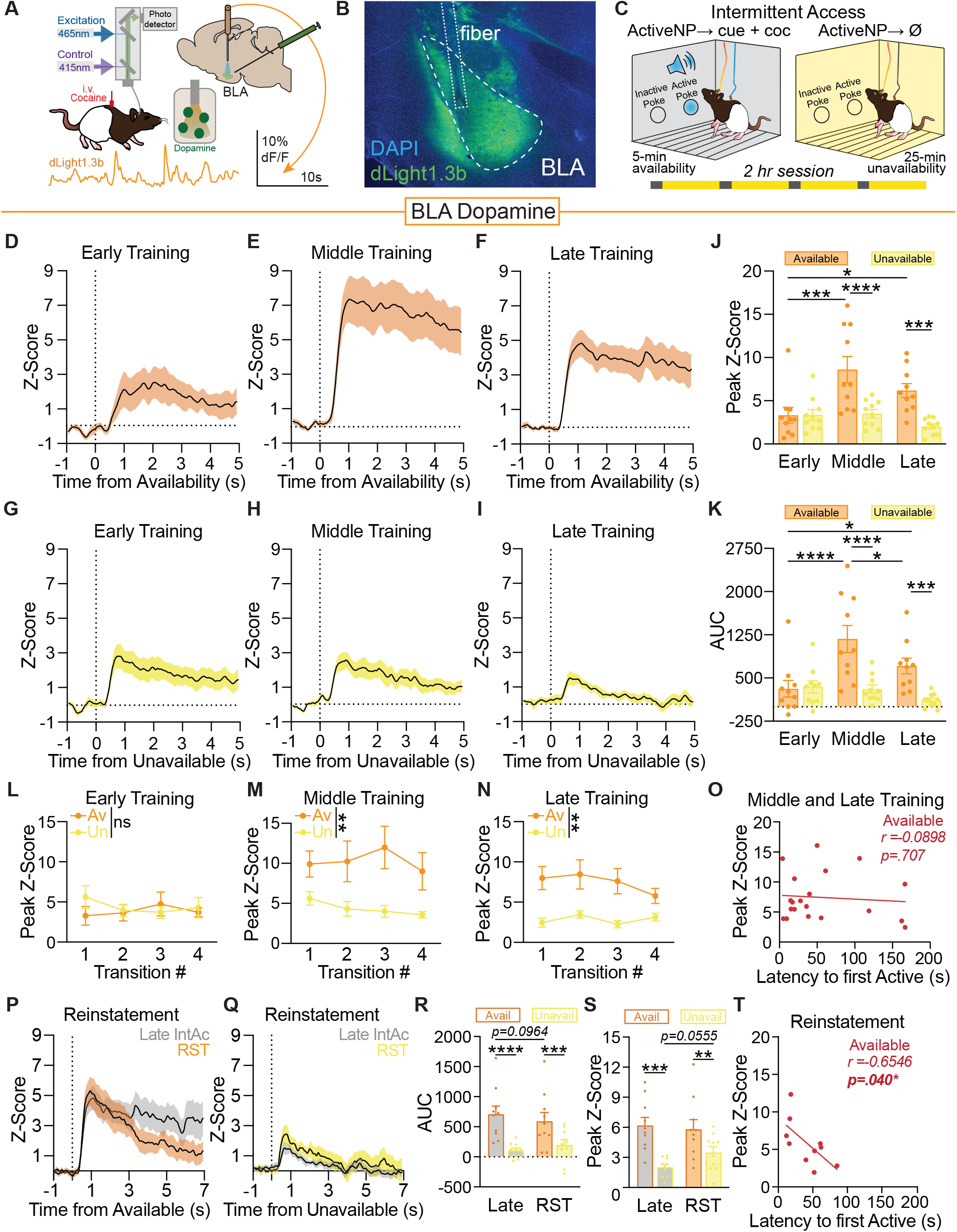
Sustained BLA dopamine signals encode drug availability state transitions ▸. A) Schematic of the fiber photometry approach for recording dopamine dynamics in the BLA (n=10, 5F, 5M). B) Representative histology image of dLight viral expression and fiber damage in the BLA. C) Data depicted here are largely from the intermittent access phase which features shifts between drug available and unavailable periods signaled by changes in sensory context. D) Positive dopamine signals are seen in the BLA during the cued drug availability transitions (house lights off) in early, E) middle and F) late intermittent access. The BLA dopamine response to the unavailability cue (house lights on) during G) early, H) middle, and I) late intermittent access show mild dopamine signals across training. J) Peak measures and K) AUC quantifications of the first 5s after state shifts of the BLA dopamine signals comparing the first, middle, and last intermittent access session. L) Within each intermittent access training session there were four drug availability and unavailability transitions. During the first session, peak measures of the dopamine traces were similar for both availability state transitions. M) In middle training and N) late training, availability cue evoked dopamine was generally greater than the peak of the unavailability cue across the individual transition trials. O) There was no correlation between the availability transition dopamine response and the latency to the first active nose poke during the middle and end of intermittent access training. P) Robust BLA dopamine signaling was observed following the drug availability period onset during reinstatement, plotted against the BLA dopamine trace recorded in late intermittent access training (gray). Q) Sparse BLA dopamine dynamics were evoked by the drug unavailability transition in conditioned reinstatement, plotted with the BLA dopamine trace from late intermittent access training (gray). The drug availability shift prompted larger dopamine release than the drug unavailability shift during both the final intermittent access session and reinstatement in terms of R) AUC quantifications of the first 5s after state shifts and S) peak measures. T) The drug availability shift related dopamine signal negatively correlated with latency to first active nose poke during the reinstatement test. Data represent mean ± SEM. *p<0.05, **p<0.01, ***p<0.001, ****p<0.0001. AUC: Area under curve. RST: reinstatement.

To explore the time course of these patterns further, we analyzed dopamine signals within sessions, isolating the 4 drug availability and 4 drug unavailability transitions in a trial-like fashion. In the first intermittent access session, there were no consistent differences in dopamine response between transition types (**Fig 3L**; mixed-effects ANOVA, no main effect of state type: F(1,9)=0.1393, p=0.7176). As training progressed, the availability transitions consistently promoted larger dopamine signals within middle training (**Fig 3M**; mixed-effects ANOVA, main effect of state type: F(1,9)=11.95, p=0.0072) and late training sessions (**Fig 3N**; mixed-effects ANOVA, main effect of state type: F(1,9)=20.89, p=0.0013). Notably, in the middle and late training phases, discrimination of availability and unavailability transitions was evident in the dopamine response at the first availability transition of the day, before any cocaine had been taken (**Fig S5**). This indicates that BLA dopamine dynamically reports learned and motivationally salient state transitions in the context of drug self administration, rather than solely tracking sensory changes or drug exposure.

We next looked at dopamine responses during the reinstatement test following extinction. Here, the reintroduction of dynamic state shifts prompted robust BLA dopamine response to the drug “availability” transition versus a small response to the drug unavailability transition (**Fig 3P-S**; two-way ANOVA, main effect of state type, peak: F(1,9)=18.53, p=0.0020; AUC: F(1,9)=13.98, p=0.0046), and the drug availability transitions evoked significantly greater dopamine compared to the unavailability (peak: p=0.0065; AUC: p=0.0002). The transition signals seen in reinstatement were similar to those from the late stage of intermittent access training, but did change subtly (**Fig 3R**; AUC: phase x state type interaction: F(1,9)=5.308, p=0.0467). While the drug availability transition elicited similar peak values of BLA dopamine in late intermittent access and reinstatement (**Fig 3S**; p=0.5661), the degree of sustained dopamine response trended lower during reinstatement (**Fig 3R**; p=0.0964). Further, evoked dopamine to the drug unavailability transition in reinstatement trended larger than in late intermittent access (**Fig 3S**; peak: p=0.0555).

Finally, we explored the relationship between state transition evoked dopamine signals and behavior. First, focusing on the middle and late phases of intermittent access training, we found that the magnitude of dopamine signals evoked by the availability transition did not correlate with rat’s latency to make their first active nose poke (**Fig 3O**; Pearson correlation, r=-0.0898, p= 0.7066). In contrast, in the reinstatement condition, we found a significant correlation, such that larger availability transition evoked dopamine signals predicted shorter latency to first drug seeking nose pokes (**Fig 3T**; peak: Pearson correlation, r=-0.6546, p= 0.0400); AUC: r=-0.743, p=0.014). Together, these data suggest that state transition evoked dopamine signals in the BLA may become decoupled from behavioral expression while cocaine is onboard during active drug self administration, and hint at a complex interaction of effects on the system following drug seeking extinction and abstinence.

### Drug availability state shifts evoke transient BLA D1-neuron activity

We next examined calcium activity in BLA D1+ neurons during the intermittent access self-administration paradigm (**Fig 4A-C**). We saw robust activity in D1 neurons in response to the drug availability state onset in early (**Fig 4D**), middle (**Fig 4E**), and late intermittent access training (**Fig 4F**). Relative to dopamine signals (**Fig 3**), D1-neuron signals were comparatively transient, returning to approximate baseline levels around 5s after the onset of drug availability (**Fig S4E-H**). The corresponding shift back to drug unavailability also prompted transient BLA D1-neuron activity across all intermittent access training (**Fig 4G-I**). However, as training progressed, D1 neuron responses came to discriminate availability and unavailability transitions, as measured by signal peak (**Fig 4J**; two-way ANOVA, session x transition interaction: F(2,16)=4.232, p=0.0335) and AUC of the first 2s after state transition (**Fig 4K**; two-way ANOVA, session x transition interaction: F(2,16)=4.808, p=0.0232). Availability transition responses were larger in middle (peak: p=0.0218; AUC: p=0.0330) and late training phases (peak: p=0.0345; AUC: p=0.0308). While the D1-neuron responses to the drug availability onset were consistent between early and late training (peak: p=0.5094; AUC: p=0.5679), the drug unavailability transition signals significantly decreased between early and late sessions (AUC: p=0.0337; peak: p=0.0843), suggesting that BLA D1-neurons may respond to state-guided drug seeking preferentially by scaling down responsivity to less salient stimuli.

**Fig 4.**
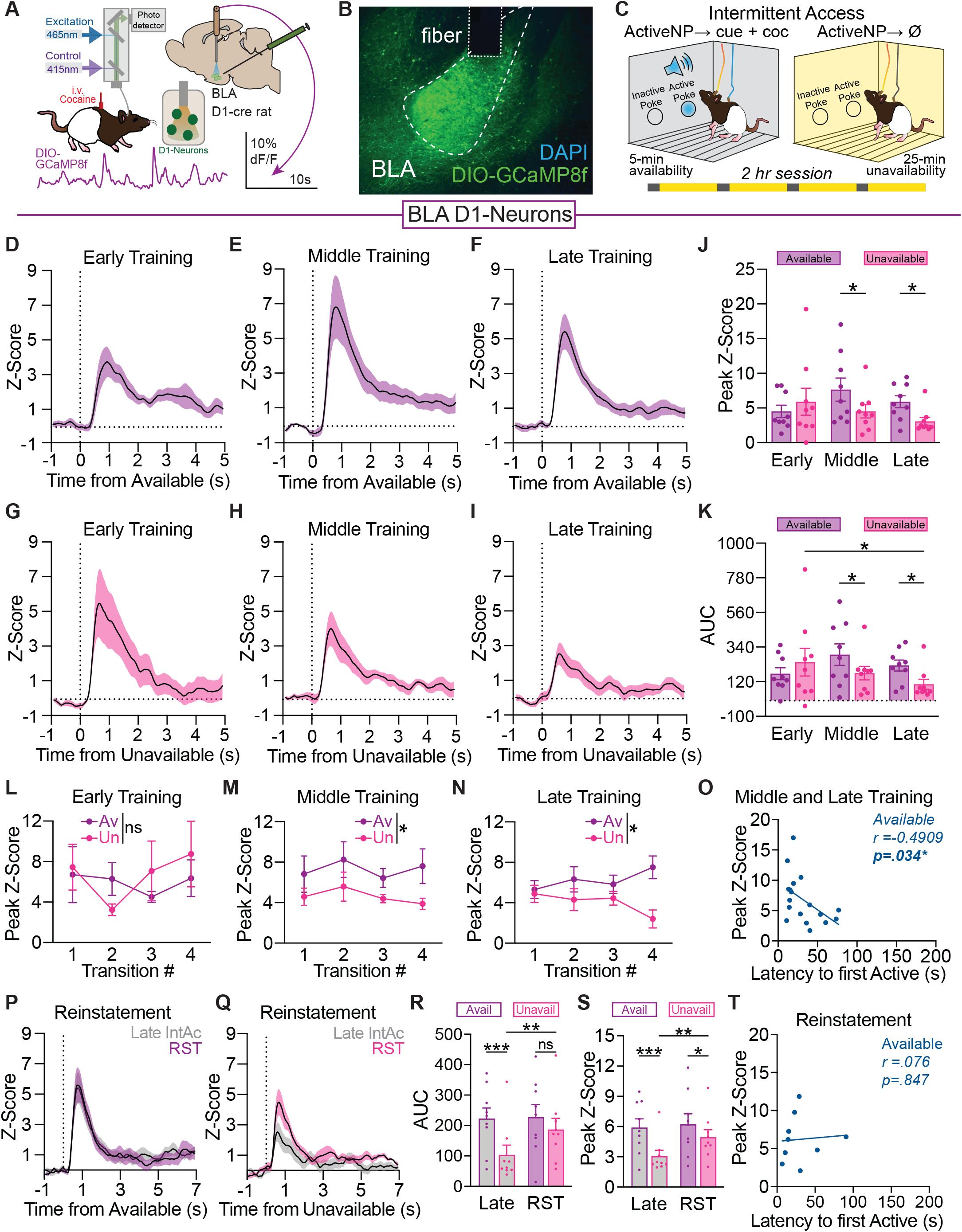
Drug availability state shifts evoke transient BLA D1-neuron activity ▸. A) Schematic of the fiber photometry approach for recording D1-neuron activity in the BLA (n=9, 5F, 4M). B) Representative histology image of cre-driven GCaMP viral expression in the D1-neurons and fiber damage in the BLA. C) Data depicted here are largely from the intermittent access phase which features shifts between drug available and unavailable periods signaled by changes in sensory context. D) The drug availability state onset (house lights off) prompted an increase in BLA D1-neuron activity in early, E) middle, and F) late intermittent access training. The BLA D1-neuron response to the unavailability cue (house lights on) across G) early, H) middle, and I) late intermittent access training sessions. J) Peak measures and K) AUC quantifications of the first 2s after the state shifts of the BLA D1-neuron activity traces comparing signals during early, middle, and late intermittent access. L) There were four drug availability and unavailability transitions in each intermittent access training session. Within the first session, D1-neuron responses to both cued availability transitions were comparable in magnitude. M) During middle training, BLA D1-neurons had larger activity following the availability cue compared to the unavailability cue throughout the intermittent access session. N) The differences in evoked activity levels were maintained in late intermittent access training. O) The D1-neuron response to drug availability was negatively correlated with latency to first active nose poke during the middle and end of intermittent access training. P) Strong activation of BLA D1-neurons were observed during cued drug availability transitions during reinstatement, plotted against the activity trace from late intermittent access training (gray). Q) Marked BLA D1-neuron dynamics were elicited by the drug unavailability cue in conditioned reinstatement, plotted with the BLA D1-neuron trace from late intermittent access training (gray). R) AUC quantifications of the first 2s after state shifts and S) peak measurements comparing reinstatement to the final intermittent access session. T) There was no correlation between latency to first seeking bout and the availability cue D1-neuron signal peak during the reinstatement test. Data represent mean ± SEM. *p<0.05, **p<0.01, ***p<0.001, ns: not significant. AUC: Area under curve. RST: reinstatement.

We next examined the within session patterns of BLA D1 neuron activity across individual transitions. In the first intermittent access session, D1-neurons did not consistently distinguish availability and unavailability transitions (**Fig 4L**; mixed-effects ANOVA, no main effect of state type: F(1,8)=0.1029, p=0.7566). As training progressed, D1-neurons preferentially responded to the availability transitions in the middle (**Fig 4M**; mixed-effects ANOVA, main effect of state type: F(1,8)=7.023, p=0.0293) and late phases (**Fig 4N**; mixed-effects ANOVA, main effect of state type: F(1,8)=7.491, p=0.0256). Within the final session, D1 neuron signals further discriminated state transition types (**Fig 4N**; trial x cue type interaction: F(3,20)=3.431, p=0.0367). Similar to dopamine signals, D1 neurons had a strong response to the first drug availability transition in middle and late training sessions (**Fig S5**), before any cocaine had been taken on the day.

We next looked at D1 responses during the reinstatement test after extinction. Reintroducing the dynamic state shifts prompted robust D1-neuron activity in response to the drug “availability” transition and a slightly smaller response to the drug unavailability transition (**Fig 4P-S**; two-way ANOVA, main effect of state type, AUC: F(1,8)=7.246, p=0.0274; peak: F(1,8)=9.309, p=0.0158). D1-neuron discrimination of availability and unavailability transitions was not as distinct in reinstatement compared to the final day of intermittent access training (**Fig 4R**; AUC: phase x state type interaction: F(1,8)=6.015, p=0.0398), where during reinstatement, both state transitions evoked comparable levels of D1-neuron activity (AUC: p=0.1155). Notably, while availability transition signals were the same in reinstatement compared to late intermittent access training, unavailability transition responses were significantly larger in reinstatement relative to late intermittent access (AUC: p=0.0062; peak: p=0.0051). The peak measure showed more consistent discrimination between states in reinstatement (**Fig 4S**; trend for phase x state type interaction, F(1,8)=5.081, p=0.0542) as availability transitions evoked larger D1-neuron responses compared to the unavailability transitions (p=0.0327).

Finally, we explored the relationship between state transition signals in D1 neurons and behavior. In contrast to the dopamine signals, during the middle and final intermittent access sessions, we found a significant negative correlation between the peak in D1-neuron activity evoked by the drug availability transition and rat’s latency to their first active nose poke (**Fig 4O**; Pearson correlation, r=-0.4909, p= 0.0386), such that larger signals predicted faster drug seeking responses. Additionally, larger D1-neuron responses were correlated with seeking behavior in minute one (Pearson correlation, r=0.5693, p= 0.0137) and the full availability period (Pearson correlation, r=0.5410, p= 0.0204). However, this correlation between D1-neuron activity and behavior was no longer present during the reinstatement test (**Fig 4T**; Pearson correlation, r=0.0757, p= 0.8465). Together, these results suggest a possible synergy between active state transition evoked drug seeking motivation and D1-neuron activity, which may be reduced when drug is no longer available.

### Drug availability states preferentially engage the BLA dopamine system and modulate drug seeking action-related signals

We next investigated BLA dopamine signaling and D1-neuron activity during drug seeking behaviors (active nose pokes) that occurred in drug availability versus unavailability windows to explore the interaction of states and actions. Overall, active nose pokes made during the availability period coincided with larger BLA dopamine responses, compared to the same action made during unavailability periods (**Fig 5A-D**; mixed-effects ANOVA, main effect of state: F(1,9)=8.821, p=0.0157). This distinction was especially pronounced in early (p=0.0013) and middle (p=0.0133) training, but diminished by late training (p=0.2661). Similarly, D1 neurons were more engaged during active nose pokes during the availability period (**Fig 5I-J**); two-way ANOVA, main effect of state: F(1,8)=10.05, p=0.0132), and this distinction trended away by late training.

**Fig 5.**
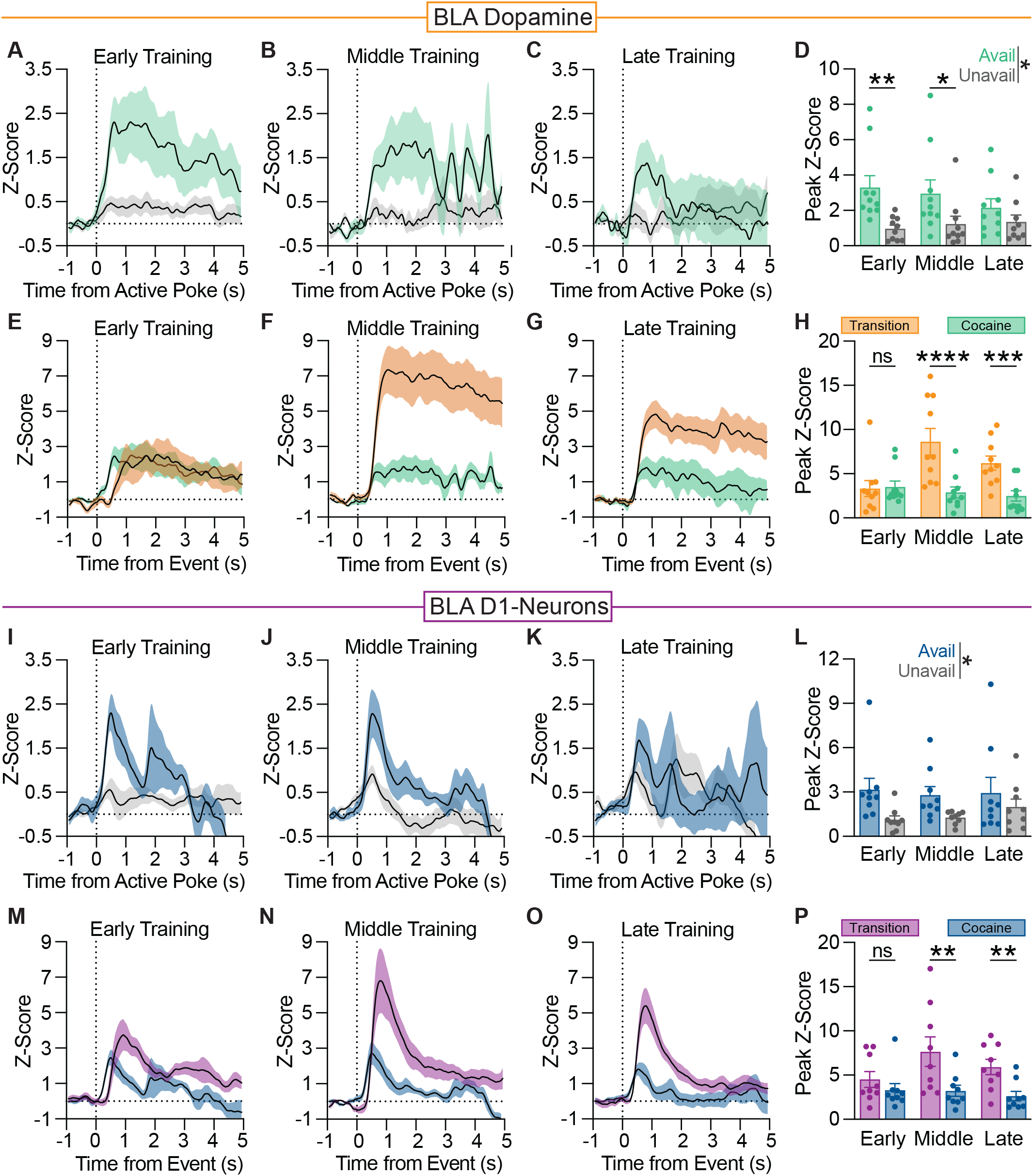
Drug availability states preferentially engage the BLA dopamine system and modulate drug seeking actionrelated signals ▸. (A)BLA dopamine (dLight: n=10) responses following all active nose pokes made within the drug available vs unavailable periods in early (B)middle, and C) late intermittent access training. D) The peak dopamine signal for active nose pokes within the availability period was larger than for those made within the unavailable period. E) BLA dopamine activity to the drug availability transition compared to proximal drug seeking actions (active nose pokes resulting in an intravenous cocaine infusion and delivery cue) in early, F) middle, and G) late in intermittent access training. H) The peak signal for availability state transitions grew with training while the cocaine peak signal did not change. I) BLA D1-neuron (GCaMP: n=9) signals following all active nose pokes made within the drug available and unavailable periods in early J) middle, and K) late intermittent access training. L) Generally, the peak D1-neuron response to active nose pokes within the availability period was larger than active nose pokes during the unavailable period across the intermittent access paradigm. M) BLA D1-neuron signals following the drug availability transition and cocaine-paired active nose pokes in across early, N) middle, and O) late intermittent access training. P) The peak signal for availability state transition grew, relative to the cocaine signal, which did not change. Data represent mean ± SEM. *p<0.05, **p<0.01, ***p<0.001, ****p<0.0001, ns: not significant.

Larger active nose poke associated photometry signals seen in the availability period during intermittent access could be related to the presentation of drug delivery cues and/or cocaine infusions. To explore this further, we next looked at active nose poke-related signals in the reinstatement test, where drug delivery cues, but no cocaine, accompanied active nose pokes in the “availability” period. During reinstatement, we no longer saw divergent encoding of active nose pokes based on the availability state they occurred in for either BLA dopamine nor D1-neuron recordings (**Fig S5E,F,M,N**). This suggests a complex interaction of drug availability state and cocaine exposure drives preferential encoding of drug seeking actions during intermittent access, and that the BLA dopamine system is relatively less reactive to more proximal drug delivery cues.

We next directly compared dopamine signals evoked by drug availability transitions and active nose pokes made in the availability window that resulted in cocaine infusions. These signals co-varied across intermittent access training (**Fig 5E-H**; two-way ANOVA, phase x event type interaction F(2,18)=10.33, p=0.0012; main effect of event, F(1,9)=25.02, p=0.0007). In early training, there was no difference in state versus action signals (p=0.8295), however by middle (p<0.0001) and late (p=0.0007) intermittent access, availability transitions evoked much larger dopamine signals compared to cocaine seeking actions. We saw a similar pattern for D1-neurons in the BLA, which were preferentially activated following drug available period transitions compared to active nose pokes resulting in cocaine infusions (**Fig 5M-P**; main effect of event, F(1,8)=6.033, p=0.0395). State transition and cocaine seeking D1-neuron responses were similar early in intermittent access training (p=0.2905), but availability transitions came to evoke much larger D1 signals in middle (p=0.0011) and late (p=0.0094) training.

This preferential state transition encoding persisted in the reinstatement test, where drug “available” transitions evoked larger dopamine and D1-neuron signals compared to active nose pokes (**Fig S5G,H,O,P**). Further, when comparing photometry signals at the first drug availability transition and first active nose poke made in each intermittent access training session, thus before any cocaine had been taken, we still saw preferential state transition activity in dopamine (**Fig S5A-D**) and D1 recordings (**Fig S5I-L**). This suggests cocaine’s pharmacological actions do not fully account for exaggerated state signals during self administration.

Collectively, these data suggest that the BLA dopamine system preferentially signals drug availability states over drug seeking actions, proximal cues, and drug infusions, particularly as behavior becomes heavily governed by sensory transitions signaling intermittent drug availability.

### Increasing BLA D1-neuron activity at drug availability transitions enhances the vigor of cocaine binging

Collectively, our data indicate that the BLA dopamine system is dynamically engaged as rats develop sensory guided drug self administration behavior. To interrogate the functional role of some of these signals, we next used brief optogenetic manipulations of BLA D1 neurons. Optogenetic excitation was delivered at the onset of each drug availability state transition (extinguished box lights) during three sessions (Avail Stim) across intermittent access training: early, middle, late (**Fig 6A**). Intermittent access sessions without any optogenetic manipulations (No Stim) within the same training phases were used as behavioral comparisons. As with our previous cohorts, rats came to discriminate between the drug available and unavailable periods, titrating behavior in a sawtooth fashion (**Fig 6B**). Total drug intake was not impacted by our focused optogenetic manipulations, as Cre+ Early Middle Late Early Middle Late Early Middle Late and Crecontrol groups took the same number of cocaine infusions (**Fig 6C**; two-way ANOVA, main effect of virus: F(1,9)=0.1750, p=0.6855; stim x virus interaction: F(1,8)=0.2421, p=0.6359). Overall, D1-cre+ rats did not take more cocaine infusions on optogenetic stimulation sessions, compared to no stimulation sessions (**Fig 6D**; two-way ANOVA, main effect of stim: F(1,5)=1.448, p=0.2827), although we saw a trend for enhanced intake in the middle training phase (p=0.0770).

**Fig 6.**
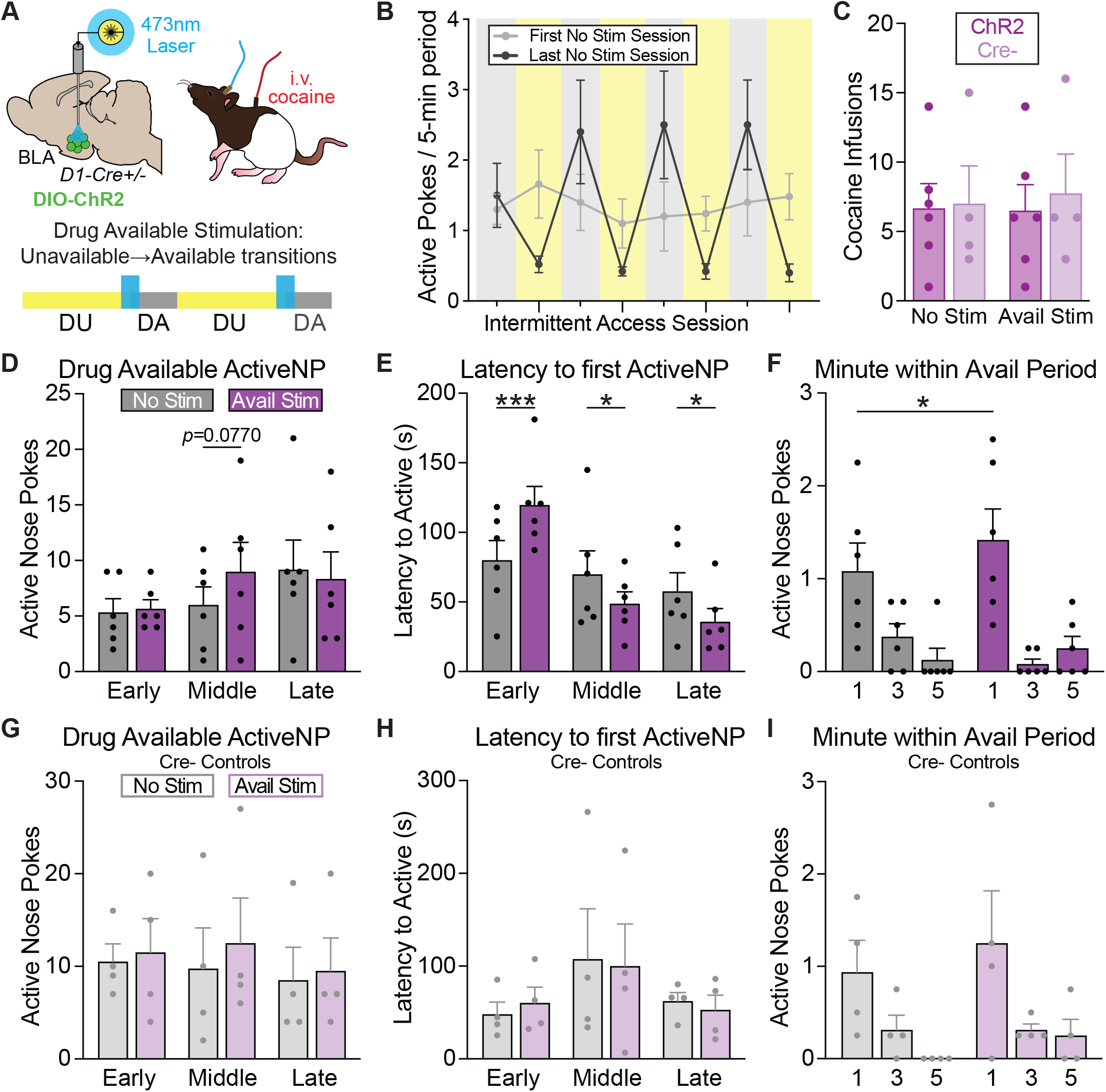
Increasing BLA D1-neuron activity at drug availability transitions enhances the vigor of cocaine binging. A) Schematic illustrating the approach to drive temporally precise optogenetic activation of BLA D1+ neurons in vivo using a 20Hz, 5s 473-nm laser pulse. Drug available manipulations were targeted to the onset of each drug availability state transition (extinguished box lights) on a subset of sessions. B) Rats (n=10, 5F, 5M) learned to discriminate between the drug available and unavailable periods during intermittent access training. C) Rats in both viral conditions (ChR2/Cre+, n=6, and Crecontrol, n=4) took similar amounts of cocaine infusions during late intermittent access training on the final no stimulation vs stimulation day. D) Average total active nose pokes earned during the drug available periods during stimulation (purple) and no stimulation (gray) sessions in early, middle, and late intermittent access training. E) Latency to the first active poke across intermittent access for the stimulation vs no stimulation conditions. BLA D1-neuron activation at drug availability onset decreased the latency of cocaine seeking compared to no stimulation sessions in middle and late training. F) Quantification of active nose pokes made in minutes 1, 3, and 5 of the drug available periods during the final no stimulation and stimulation sessions. Active pokes in the 1st minute of the stimulation session were elevated compared to the no stimulation session. G) Creviral controls underwent the same intermittent access training and laser delivery schedule. Active pokes during the drug available periods between stimulation conditions across intermittent access training in the viral controls. H) Latency to the first active poke of the viral control group across training progression and stimulation conditions. I) Invigoration of the binge-like bouts in the first minute of the drug availability period was not impacted by laser delivery in controls. Data represent mean ± SEM. *p<0.05, ***p<0.001. DA: drug available. DU: drug unavailable. NP: nose poke.

Amplifying BLA D1-neuron activity impacted the latency of drug-seeking responses during availability periods as intermittent access training progressed (**Fig 6E**; twoway ANOVA, stim x session interaction: F(2,10)=17.62, p=0.0005). D1 neuron excitation significantly reduced the latency of drug-seeking responses in both middle (p=0.0302) and late (p=0.0271) intermittent access training (**Fig 6E**). Again, rats developed binge-like seeking patterns where the distribution of active nose pokes were concentrated to the first minute of drug availability (**Fig 6F**; two-way ANOVA, main effect of minute: F(2,10)=13.13, p=0.0016). In late training, excitation of BLA D1-neurons invigorated this binge-like seeking, such that more active nose pokes were concentrated in minute 1 of drug availability compared to No Stim sessions (**Fig 6F**; two-way ANOVA, minute x stim interaction: F(2,10)=11.36, p=0.0027; Avail Stim vs. No Stim: p=0.0458). We saw no effect of laser delivery on behavior in Cre-control rats on total drug seeking actions (**Fig 6G**; two-way ANOVA, main effect of stim: F(1,3)=6.987, p=0.0774), latency to first nose poke (**Fig 6H**; two-way ANOVA, main effect of stim: F(1,3)=0.09877, p=0.7739), and minute 1 seeking invigoration (**Fig 6I**; two-way ANOVA, minute x stim interaction: F(2,6)=0.5294, p=0.6141). These results indicate that transient excitation of BLA D1 neurons is important for enhancing the vigor of cocaine seeking bouts once behavior is under the control of dynamic environmental state shifts.

### Inhibition of BLA D1 neurons at drug availability transitions does not impact the magnitude or timing of binge-like cocaine seeking

In a final set of studies, we used brief optogenetic inhibition to explore whether activation of BLA D1-neuron activity during state transitions is necessary to promote cocaine binging (**Fig 7A**). Optogenetic inhibition paired with a subset of availability transitions early, middle, and late in intermittent access did not affect acquisition of the typical saw-tooth cocaine seeking pattern (**Fig 7B**). Rats in both the Cre+ and Cre- control viral groups earned similar amounts of cocaine during late intermittent access on both stimulation and non-stimulation sessions (**Fig 7C**; two-way ANOVA, main effect of virus: F(1,12)=0.5044, p=0.4912; stim x virus interaction: F(1,12)=0.8865, p=0.3650).

**Fig 7.**
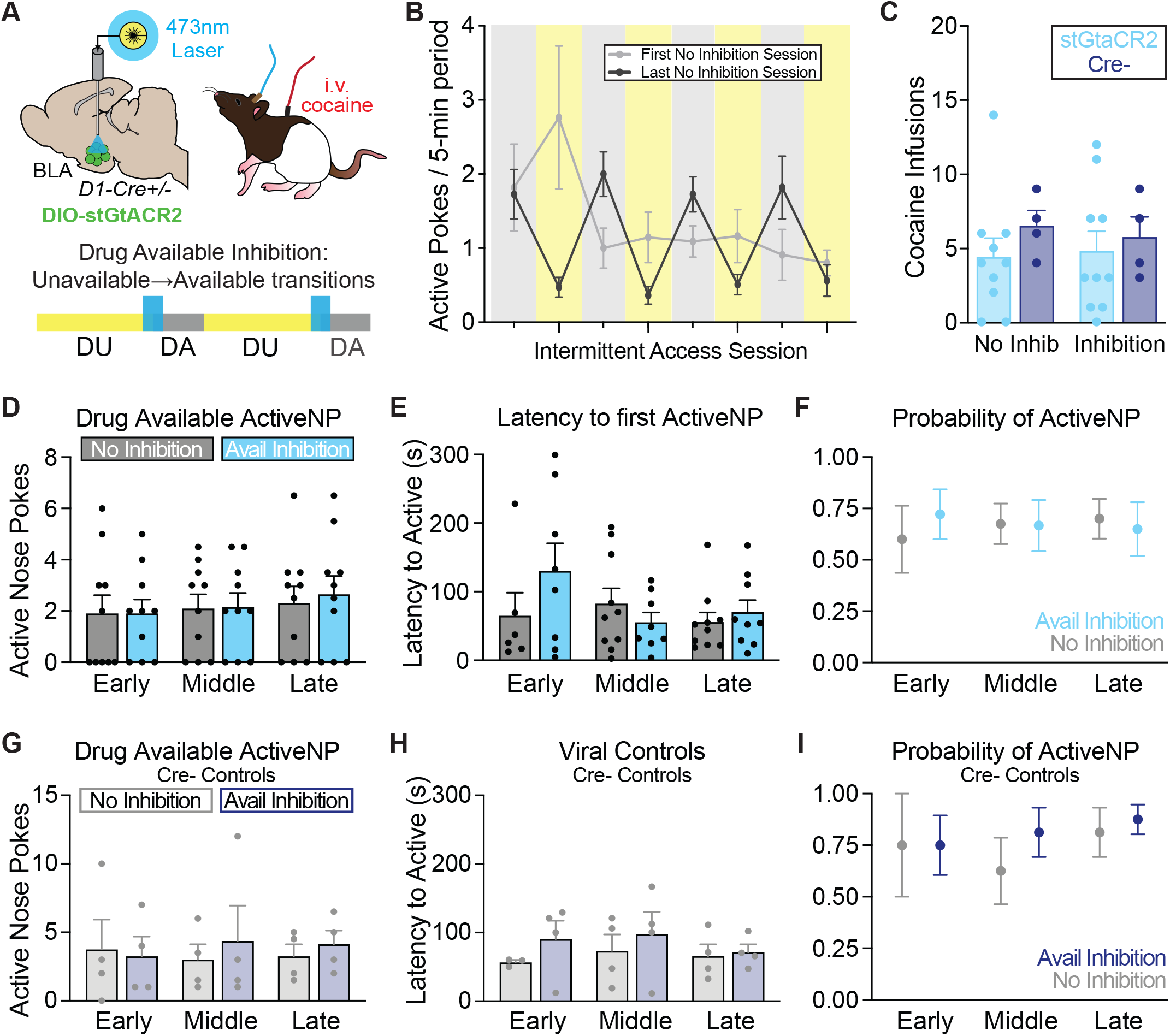
Silencing BLA D1-neuron activity during drug availability transitions does not impact cocaine seeking. A) Viral approach to drive temporally precise silencing of BLA D1+ neurons in vivo. Manipulations were targeted to the onset of the drug availability state (extinguished box lights) using a constant 3s 473-nm laser pulse. During inhibition sessions, half of the availability transitions were paired with optogenetic inactivation. B) Behavior data comparing the first and last cocaine intermittent access training sessions without optogenetic manipulations (day 1 and day 14). Rats (n=14, 9F, 5M) learned to discriminate between the drug available and unavailable periods with training. C) Rats in both viral conditions (stGtaCR2/Cre+, n=10, and Crecontrol, n=4) took a similar number of cocaine infusions during the final no inhibition vs inhibition days. D) Average total active nose pokes during the drug available periods during inhibition vs no inhibition trials in early, middle, and late intermittent access training. E) Latency to the first active poke across training and inhibition conditions. F) Probability of an active poke event occurring during the drug available periods throughout intermittent access training. G) Viral controls underwent the same intermittent access training paradigm and laser delivery schedule. Active nose pokes during the drug available periods between inhibition conditions across intermittent access training for the control group. H) Latency to the first active poke of the viral control group across training progression and stimulation conditions. I) Probability of an active poke occurring during the drug available periods throughout intermittent access training for the viral controls. Data represent mean ± SEM. DA: drug available. DU: drug unavailable. NP: nose poke.

We found no effect of transient BLA D1-neuron inactivation at availability transition onsets on the overall amount of active nose pokes made compared to no laser trials (**Fig 7D**; mixed-effects ANOVA, no main effect of stim: F(1,9)=0.9351, p=0.3588). Further, optogenetic inhibition did not delay the initiation of drug-seeking responses during intermittent access (**Fig 7E**; two-way ANOVA, main effect of stim: F(1,9)=0.7408, p=0.4117). Finally, inactivation of BLA D1-neurons did not affect the probability of a drug seeking event during the drug availability periods (**Fig 7F**; two-way ANOVA main effect of stim: F(1,9)=0.04997, p=0.8281). We similarly saw no effect of laser delivery on behavior in Cre- controls (**Fig 7G-I**; two-way ANOVA, total nose pokes: F(1,3)=3.267, p=0.1684; latency to first poke: F(1,3)=3.245, p=0.1695; probability of a poke F(1,3)=1.500, p=0.3081).

Together, results from our bidirectional optogenetic manipulations (**Fig 6,7**) indicate that brief, phasic activity in BLA D1+ neurons does not gate the expression of state-guided drug seeking, but is permissive for vigorous sensory guided drug seeking motivation.

## DISCUSSION

We examined how the BLA dopamine system is engaged during the development of dynamic, sensory-guided drug seeking behavior. First, we show that cocaine exposure sensitizes dopamine transmission from VTA dopamine neurons projecting to the BLA. Second, we used an intermittent access cocaine self-administration paradigm that prompted robust binge-like cocaine seeking heavily guided by shifts in the sensory environment. The transitions to drug available states evoked sustained BLA dopamine signals as well as transient calcium responses in BLA D1+ neurons. Third, BLA dopamine signals and D1-neuron activity came to preferentially encode these drug availability states over unavailability states and relative to drug seeking actions. Dopamine signals were predictive of drug seeking behavior in a cocaine-free relapse test, while D1-neuron signals predicted drug seeking during self administration training. Finally, we found that brief optogenetic enhancement of BLA D1-neuron activity during drug availability state transitions potentiated the vigor of cocaine binging, while optogenetic inhibition did not affect behavior. Collectively, our studies pinpoint the dopaminergic amygdala as a key interface between the environment and decisions to engage in vigorous drug use. These results support an addiction framework where drug seeking motivation is dynamically controlled by hierarchically organized environmental stimuli that are encoded by dopamine signaling and neuronal ensembles within the amygdala (Paton et al., 2006; Tye and Janak, 2007; Belova et al., 2008; Gründemann and Lüthi, 2015; Saez et al., 2015; Wassum and Izquierdo, 2015; O’Neill et al., 2018; Sengupta et al., 2018; Zhang and Li, 2018; Gründemann et al., 2019; Gostolupce et al., 2021; Zhang et al., 2021, 2025; Ottenheimer et al., 2024; Kong et al., 2025; Brickner et al., 2026).

### VTA to BLA dopamine circuitry is modulated by cocaine exposure

We found that cocaine sensitization potentiates dopamine transmission in the VTA to BLA circuit, extending past microdialysis studies showing that BLA dopamine levels increase following cocaine infusions (Hadfield and Milio, 1992; Wilson et al., 1994; Hurd et al., 1997). Given this work and our past studies (Brickner et al., 2026), we hypothesize that the VTA is likely the main source of reward and context-related dopamine signals into the BLA. Consistent with this, tonic VTA to BLA dopamine transmission contributes to drug self-administration acquisition (Smith and Torregrossa, 2024) and optogenetic excitation of VTA-BLA dopamine neurons can drive certain forms of reward learning (Sias et al., 2024). Notably, the effects of cocaine sensitization were specific to enhancing dopamine evoked by optogenetic stimulation of the VTA-BLA pathway, and did not influence dopamine evoked by an aversive stimulus. This may suggest that additional mechanisms contribute to aversion-related dopamine release, which could arrive from the periaqueductal gray and locus coeruleus (Loughlin and Fallon, 1983; Poulin et al., 2018; Zafiri and Duvarci, 2022; Zhao et al., 2022; Duvarci, 2024; Matarasso et al., 2026). The impact of drug use on valence processing in the amygdala will be important to further understand from a translational perspective, given that addiction and PTSD can be comorbid conditions (McCauley et al., 2012; Tipps et al., 2014; Gilpin and Weiner, 2017; Najavits et al., 2020; Renaud et al., 2021; Hinckley and Danielson, 2022).

The brain-wide dopamine system has been heavily implicated in drug relapse (Weiss et al., 2000; Saunders et al., 2013; Liu et al., 2020; Luján et al., 2023), including dopamine input into the BLA (Tran-Nguyen et al., 1998; Torregrossa and Kalivas, 2008; Smith and Torregrossa, 2024). Cocaine exposure increases VTA dopamine neuron activity (Creed et al., 2016; Arencibia-Albite et al., 2017; Francis et al., 2019), however, chronic cocaine use can trigger the striatum into a hypodopaminergic state associated with reductions in VTA-stimulated nucleus accumbens dopamine release (Mateo et al., 2005; Siciliano et al., 2015; Francis et al., 2019; Samaha et al., 2021), so the interaction of these variables requires further study. Here, we saw augmentation of the VTA-BLA dopamine circuit specifically after a period of incubation, which suggests that an overactive BLA dopamine system could contribute to relapse susceptibility following abstinence. Our data support this, as we saw a correlation drug state-evoked dopamine signals and the intensity of drug seeking behavior in a reinstatement test.

### The basolateral amygdala dopamine system tracks dynamic drug seeking states

While microdialysis studies show BLA dopamine levels steadily increase during reward seeking and drug taking (Wilson et al., 1994; Hurd et al., 1997), the dynamic encoding of drug-related states and actions we report here cement the notion that this system is broadly engaged to signal emotional salience (Lutas et al., 2019; Tang et al., 2020; Brickner et al., 2026). Further, our BLA D1+ neuron recordings expand on past electrophysiological studies during drug-related learning (Neisewander et al., 2000; Carelli et al., 2003), demonstrating an amygdalar cell type that encodes shifts in drug seeking behavior. We show transitions to drug availability evoke sustained dopamine signals in the BLA, which may reflect the enhanced emotional salience of that sensory shift (Brickner et al., 2026). Critically, this sustained pattern was evident on the first drug availability transition before drug seeking had begun on a given session, indicating that it was not related to cocaine’s actions on the dopamine transporter (DAT). Notably, the amygdala has lower levels of the DAT compared to the striatum (Revay et al., 1996; Holleran et al., 2020). This results in slower dopamine clearance and likely sets the stage for the sustained cue-evoked dopamine we see as well as the ability for incoming dopamine to prime D1-neurons to promote state-guided behaviors. This interpretation is supported by the more transient nature of the D1 neuron responses we report and the correlation between these signals and the vigor of cocaine seeking. Given cocaine’s actions to block DAT, this positions the BLA as a site for unique drug-related plasticity compared to other dopamine terminal fields. This is consistent with past studies showing that pharmacological blockade of DAT in the BLA engages reward learning-related plasticity via D1 receptors (Tye et al., 2010). Further, the slower timescale dopamine signals supported by the BLA’s cellular and circuit architecture may facilitate its role in state level, model-based, or hierarchical learning processes, compared to canonical reward prediction error or reinforcement systems (Belova et al., 2008; Prévost et al., 2013; Saez et al., 2015; Gostolupce et al., 2021; Fraser and Janak, 2023; Aguirre et al., 2024; Sias et al., 2024; Brickner et al., 2026). Model-based decision making is known to be disrupted in addiction-related behavior models, and our results highlight the BLA dopamine system as a promising target to investigate this pathology (Wied et al., 2013; Groman et al., 2019).

Within the BLA, dopamine terminals form a majority of their synapses onto BLA projection neurons (Muller et al., 2009), suggesting that our effects of D1 neuron modulation are likely due to actions downstream or parallel circuitry in the amygdala or elsewhere (Setlow et al., 2002; Di Ciano and Everitt, 2004; Ambroggi et al., 2008; Stuber et al., 2011; Namburi et al., 2015; Burgos-Robles et al., 2017; Baimel et al., 2019; Puaud et al., 2021). Additionally, while much sparser, dopamine D2 receptors are also expressed in the BLA (Greba et al. 2001; Guarraci et al. 2000; Nader & LeDoux 1999; Ponnusamy et al. 2005; Uhlén et al. 2015), but have been more directly implicated in the interaction of valenced stimuli and drug-related behaviors within the central amygdala (Thiel et al., 2010; Kim et al., 2018; Casey et al., 2022; Bijoch et al., 2025; Jo et al., 2026).

### BLA control of sensory-guided drug seeking

Our bidirectional optogenetic experiments provide evidence that functional recruitment of BLA D1-neurons during drug availability state transitions allows sensory information to dynamically drive cocaine seeking patterns. These findings contextualize past pharmacological studies implicating activation of D1 receptors in the BLA in the output of cue-driven behaviors related to natural and drug rewards (See et al., 2001; Berglind et al., 2006; Mashhoon et al., 2009; Tye et al., 2010; Ball et al., 2011; Ng et al., 2018; Kim and Lattal, 2019; Rezaei et al., 2022). Here, phasic activation D1-neurons potentiated the vigor of binging but did not increase overall cocaine intake, suggesting this population contributes to the initiation of drug-seeking actions, potentially via the encoding of environmental state information (Sengupta et al., 2018; Tang et al., 2020; Fustiñana et al., 2021). In contrast, we did not see an effect of optogenetic inhibition of D1+ neurons in the BLA on drug seeking. Our manipulations were brief, by design, and it remains possible that more chronic BLA D1 inhibition could reveal different functional roles on slower timescales. It is also likely that redundant systems responsive to drug-associated sensory information, such as dopamine and glutamate inputs to the ventral striatum (Burns et al., 1994; Horvitz, 2002; Kalivas et al., 2009; Saunders et al., 2013; Yager et al., 2015), can compensate for the brief BLA D1 neuron inhibition in our studies, to support drug seeking.

Pharmacological enhancement of BLA dopamine levels potentiates learning of stimulus-reward associations and relapse behaviors during cue-maintained reinstatement (Hitchcott et al., 1997; Ledford et al., 2003), while chemogenetic inhibition of VTA dopamine input into the BLA impairs cocaine self administration (Smith and Torregrossa, 2024). This suggests that the ability of cue to regulate drug seeking requires stable tonic dopamine levels in the BLA. It remains unclear what behavioral role BLA dopamine has for sensory-guided drug seeking on shorter phasic timescales like in our studies. Critically, dopamine does not directly activate BLA D1-containing neurons, but rather acts as a neuromodulator to enhance their excitability (Scibilia et al., 1992; Rosenkranz and Grace, 1999; Kröner et al., 2005; Takahashi et al., 2010; Lutas et al., 2022). Given the number of glutamatergic BLA inputs from thalamus, cortex, and other regions (Janak and Tye, 2015; Correia and Goosens, 2016; O’Neill et al., 2018), it will be important to investigate the role of other afferents in binge cocaine seeking and their influence on D1-neurons and BLA microcircuitry. Cocaine self-administration may recruit glutamatergic inputs that prompt a unique change in D1-neuronal activity that is dissociated from dopamine signals, for example. This interpretation is supported by our data here, where dopamine signals grew in response to drug availability transitions, but D1 neuron signals were relatively stable across training. Collectively, this suggests a complex interplay among BLA inputs to control D1-neuron activity during behavior.

In summary, we demonstrate that the dopaminergic amygdala is a focal point in the brain for environmental control of the pattern and timing of drug seeking motivation. Given that decisions to engage in drug use among humans are heavily influenced by sensory cues, our results highlight potential therapeutic targets for addiction treatments.

## Funding and Acknowledgements

The authors have no biomedical financial interest or potential conflicts of interest. This work was supported by NIH grants F31 DA060004 (MAB), P30 DA048742 (MJT), and R00 DA042895, R01 MH129370, R01 MH129320, and R01 DA057292 (BTS). We thank all members of the Thomas and Saunders labs for technical support and feedback.

## METHODS

### Subjects

We used rats (N=49, 26 female, 23 male), bred on a Long Evans background. Cocaine sensitization studies (n=6, 2F, 4M) made use of transgenic tyrosine hydroxylase (TH)-cre+ rats for targeting optogenetic stimulation to VTA dopamine neurons (Witten et al., 2011). Wild type rats were used for BLA dopamine recordings (n=10, 5F, 5M). Transgenic D1-cre+ rats (Pettibone et al., 2019) were used to record BLA D1 dopamine receptor (D1DR)-containing neurons (n=9, 5F, 4M). Optogenetic experiments used separate groups of D1-cre rats for inhibitory (n=14, 9F, 5M) or excitatory (n=10, 5F, 5M) manipulations. All subjects were between 250-350g at the time of surgery and 4-9 months old during experimentation. Rats were dual-housed until jugular vein catheterization, when applicable, with littermates in a temperature and humidity-controlled colony room on a 12-hour light/dark cycle with ad libitum access to food and water. All experimental procedures were approved by the Institutional Animal Care and Use Committee at the University of Minnesota and were carried out in accordance with the guidelines on animal care and use of the National Institutes of Health of the United States.

### Viral Vectors

For fiber photometry experiments, stereotaxic surgery was used to infuse viruses (-2.7 AP, ±4.9 ML, -8.2 DV) and implant the optic fibers (-2.7 AP, ±4.9 ML, -8.0 DV) in the BLA. dLight1.3b (500 nL, AAV5-hSyn-dLight1.3b, University of Minnesota Viral Vector and Cloning Core) (Patriarchi et al., 2020) was used to measure BLA dopamine dynamics. GCaMP8f (500 nL, AAV5--syn-FLEX-jGCaMP8f-WPRE, Addgene) (Zhang et al., 2023) was used to measure BLA D1-neuron calcium activity. For inhibitory optogenetics experiments, a cre-dependent inhibitory opsin (500nL/hemisphere, AAV1-hSyn1-SIO-stGtACR2-FusionRed, Addgene) (Mahn et al., 2018) was injected bilaterally into the BLA. For excitatory optogenetics experiments, a cre-dependent excitatory opsin (500nL, AAV5-Ef1a-DIO-hChR2(E123A)-EYFP, Addgene) virus was injected unilaterally in the BLA. For control groups, the cre-dependent virus was injected into D1-cre negative rats and the lack of viral expression was confirmed with fluorescent microscopy (**Fig S1**). To simultaneously record dopamine signals in the BLA while exciting VTA dopamine cell bodies, we injected TH-cre positive rats with a cre-dependent, excitatory, red-shifted opsin, Chrimson (1.0μL, AAV5-Syn-FLEXChrimsonR-tdTomato, Addgene) (Klapoetke et al., 2014) unilaterally into the VTA (-5.6 AP, ± 0.7 ML, -8.0 DV) and an optic fiber was placed over the VTA (-5.6 AP, ± 0.7 ML, -7.9 DV). dLight1.3b injections and optic fiber implants were targeted to the ipsilateral BLA.

### Surgical Procedures

Viral surgery: Rats were anesthetized with 5% isoflurane and placed in a stereotaxic frame, after which anesthesia was maintained at 1-3%. Saline (0.5 mL), carprofen anesthetic (5 mg/kg), and cefazolin antibiotic (70 mg/ kg) were administered subcutaneously prior to start of surgery. The top of the skull was exposed and holes were made for viral infusion needles, optic fiber implants, and 4 skull screws. A microsyringe pump delivered the viral injections at a rate of 0.1 μl/min. Injectors were left in place for 1 min, then raised 100 μm dorsal to the injection site and left in place for another 10 min to allow for diffusion, then removed slowly. Optic fibers were then chronically implanted above the injection sites and secured with skull screws and dental acrylic (Lang Dental). At the end of all surgeries, topical anesthetic and antibiotic ointment were applied to the surgical site, rats were placed on a heating pad and monitored until they were ambulatory. After stereotaxic surgery rats were paired housed with ad libitum access to food and water, given carprofen (5 mg/kg) and cefazolin (70 mg/ kg) for the first 3 days following surgery and weighed and monitored for a full week. Photometry recordings began at least 4 weeks after surgery. Optogenetic manipulations started at least 6 weeks after surgery to allow for stable virus expression.

#### Intravenous catheter surgery

Following at least 3 weeks after viral surgeries, rats received a second surgery for intra-jugular catheter placement as previously described (Collins et al., 2023). Catheters were assembled from silastic tubing (0.51mm inner diameter, 0.94mm outer diameter) epoxied to a prefabricated catheter back port (Instech). The tubing was cut to 11.5cm long and a silicone ball was constructed around the tube 2.5cm above the beveled end. For surgery, rats were anesthetized with 5% isoflurane for induction, and then maintained at 1–3%. Rats were then administered saline, carprofen anesthetic, and cefazolin antibiotic subcutaneously. Incisions were made into the skin on the back and right chest of the rat. The catheter port was affixed under the skin on the back of the rat, and the silastic tube was fed under the skin around the shoulder and into the right jugular vein. The vein was exposed and perforated to allow the tubing to be placed ∼2.5cm above the heart. Internal sutures were used to secure the silicone ball to the vein and secure the catheter in place. Both incisions were then closed with sutures and topical wound glue, and the back port was protected with a fitted cap (Instech). A topical anesthetic and antibiotic ointment were applied to the surgical sites, and rats were monitored until they were ambulatory. After this surgery rats were individually housed to protect the catheter and back port, with ad libitum access to food and water, and post-operative care as above. Catheters were flushed daily with gentamicin antibiotic (0.15 mL; 5 mg/mL) throughout recovery and the entire behavioral paradigm, to maintain patency. Twenty-four hours before experiments began, catheter patency was validated with brevital sodium (1 mg/0.1 mL, intravenous). Patency tests were repeated weekly throughout the study, and only patent rats, defined by ataxic response 3-5 s postinjection, were included in the analyses.

### Combined optogenetic stimulation-fiber photometry recordings

To gauge how dopamine dynamics in the BLA were impacted by cocaine exposure, in vivo optogenetics was paired with fiber photometry in TH-Cre rats (n=6; 2F, 4M). BLA dopamine dynamics were recorded with dLight1.3b following VTA dopamine neuron cell body stimulation using 589 nm lasers (OptoEngine and Dragon Lasers). Rats were tethered to two optic cables; one connected to the BLA implant for photometry recordings and to the VTA implant for optogenetic stimulations. Laser deliveries were not paired with any external stimulus.

#### Cocaine sensitization

A recording was collected before any cocaine was administered (pre-cocaine). Rats were first presented with 10 laser stimulations (20Hz, 2s) on a 120s fixed schedule to establish baseline evoked dopamine signals. Following this, the optic cable to the VTAwas removed and the rats experienced 5 inescapable footshocks (0.5s, 0.4mA) delivered on a 210s average variable time schedule (range: 180s-240s). The next day, rats began a cocaine sensitization paradigm, where an intraperitoneal injection of cocaine (20 mg/kg/injection) was administered approximately 24 hours apart for 4 consecutive days. 24 hours following the final cocaine injection, rats underwent a fiber photometry recording session where 10 laser stimulations and 5 footshocks were delivered with the same parameters as the pre-cocaine session. Rats then returned to their home cage and a 7 day abstinence period began. Dopamine dynamics to the laser stimulations and footshocks were recorded a final time during a 7 day post-cocaine test.

### Intermittent access cocaine self-administration paradigm

One week after catheter placement, rats began cocaine self administration using procedures modified from past studies (Calipari et al., 2013; Kawa et al., 2019; Collins et al., 2023).

#### Acquisition

First, rats learned to self-administer cocaine in 5 or 6 continuous-access sessions conducted with no ambient lighting in the chambers (house lights off). On a fixed-ratio 1 schedule of reinforcement (FR-1), active nose pokes resulted in a 20s drug-delivery cue (a tone and the illumination of the active nose port) and a delivery of a 3s intravenous injection of cocaine (0.4 mg/kg/injection). Rats could only receive one cocaine infusion per 20s cue presentation and had to wait until cue cessation before receiving another infusion. Nose pokes in the counterbalanced inactive port were recorded but had no consequences. Each session lasted 2 hours.

#### Intermittent Access training

Rats then received fourteen 2-hour daily sessions of intermittent cocaine access. Within each session, the 5-minute drug available periods were signaled by an availability signal (extinguished house lights), wherein active nose pokes resulted in the delivery of an intravenous cocaine infusion and the coincident drug-delivery cue (3s). No additional infusions could be earned during this 3s window but there was otherwise no timeout period. Following each drug availability period, the house lights reilluminated, signaling a 25-minute drug-free period where nosepokes were inconsequential. Each intermittent access session had 4 cycles of drug availability and unavailability states.

#### Extinction

Following the last intermittent access session, rats underwent 5 sessions of extinction training to reduce seeking behavior in the drug-associated state (box lights off). Each session lasted 1 hour and included one transition to drug availability (chambers lights turn off when the session began). Here, neither active nor inactive nose pokes had any programmed consequences (no cue nor cocaine deliveries).

#### Conditioned reinstatement

rats ended their training with a 2-hour reinstatement test to examine the influence reintroduction of the dynamic drug “availability” cycles (5-minute lights off, 25-minute lights on). During reinstatement, the proximal delivery cues were also present, where each active nose poke during the drug “availability” state produced the drug delivery cue (20s), but no cocaine was delivered.

### In vivo fiber photometry recordings

In vivo fiber photometry was used to measure either BLA dopamine signaling (n=16, 7F, 9M) or BLA D1+ neural activity (n=9, 5F, 4M), using established workflows (Engel et al., 2024; Stelzner et al., 2025; Brickner et al., 2026). The fiber photometry system contained optical components from Doric Lenses (Québec, Canada). This included fluorescence mini-cubes to transmit concurrent light streams from LEDs, 465nm and 405/415nm, that were sinusoidally modulated at 211Hz and 330Hz, respectively. The 465nm channel stimulated fluorescence from the dopamine-dependent dLight or the calcium-dependent GCaMP. To control for optical noise and movement artifacts, we include an isosbestic control signal (405/415nm) that is both dopamine- and calcium-independent, eliminating the need for GFP control groups. Demodulating the brightness produced by both channels allows for reliable correction of movement or photobleaching artifacts. Before each photometry recording both LED currents were set to ∼50μW. Fluorescence was transmitted from the fiber tip over the BLA back to the mini-cube via an optic cable, where it was passed through a GFP emission filter, amplified and directed into a high sensitivity photoreceiver (Newport, Model 2151) for signal processing. Meanwhile, the session was controlled and data was acquired via Synapse software running on a Tucker-Davis Technologies (TDT, Alachua, FL) real-time processor (RZ5P or RZ10X). Synapse software on the processor calibrated the output of each LED channel and simultaneously recorded photometry signals, which were being sampled from the photodetector at kHz. TTL signals from the behavioral chambers (Med Associates) were sent to the RZ processor and timestamped for post-hoc analysis. This allowed for the alignment of the real-time neural signals to behavioral (e.g., nose pokes) and environmental (e.g., cue presentations) occurrences. Video recordings were also taken and integrated with the photometry and behavior data via the Synapse software.

### In vivo optogenetic manipulations

Excitation or inhibition of D1DR-containing neurons was achieved via implantation of optic fiber in the BLA of D1-Cre rats (Pettibone et al., 2019). Cre-dependent opsins were also injected (coordinates above) to deliver the excitatory channelrhodopsin (ChR2) or an inhibitory, blue-light sensitive opsin (stGtACR2). To control for nonspecific behavioral effects of optogenetic stimulation and laser light delivery to brain tissue, experiments incorporated unpaired stimulation and Cre- control groups. Blue 473nm lasers (Dragon Lasers) were used to either excite or inhibit BLA D1-neurons. Lasers were adjusted to read ∼10mW from the end of the patch cable at constant illumination. Light output during individual 5ms light pulses during experiments was estimated to be ∼2mW/mm2 at the tip of the intracranial fiber. Light power was measured before and after every recording session to ensure that all equipment was functioning properly. Optic tethers connecting rats to the rotary joint were sheathed in a lightweight armored jacket to prevent cable breakage and block visible light transmission. During experimentation, the optic tether was connected to the optic implant via a metal sleeve to further block visible light from escaping.

#### Excitation experiments

Laser (5s, 20Hz, 10mW) was delivered at the onset of each transition to drug availability on intermittent access training days 4, 8, and 11 (Drug Available sessions). As a within-subjects control, rats received laser stimulations delivered at the transition to every drug unavailable period on intermittent access training days 3 and 12 (Drug Unavailable sessions). The remaining intermittent access training days had no optogenetic manipulations (No Stim sessions). During the reinstatement test, all of the transitions to the reintroduced drug availability were paired with optogenetic excitation delivered at cue onset.

#### Inhibition experiments

Laser (3s, continuous, 10mW) delivery began 1s before the transition to drug availability on intermittent access training days 4, 7, 10, 13, and 16 (Drug Available sessions). Early training was defined as session 4, while middle training data contained the averaged behavior in sessions 7 and 10 and late training contained the averaged behavior in sessions 13 and 16. As a within-subjects control, rats received the same optogenetic inhibition but delivered time locked the transition to drug free periods on intermittent access training days 3, 6, 9, 12, and 15 (Drug Unavailable sessions). During both types of optogenetic-paired sessions, two out of the four state transitions were paired with inhibition. The inhibition trials were pseudorandomly delivered interwoven with the non-stimulation trials. The remaining intermittent access training days had no optogenetic manipulations (No Inhibition sessions). During the reinstatement test, half of the transitions to the reintroduced drug availability were paired with optogenetic inhibition.

### Behavioral analysis

Behavioral measures (e.g., nose pokes) were automatically recorded by Med Associates software. Further analysis was analyzed using custom MATLAB pipelines to collect measures of behavior latency, rate of seeking, and binning behavior types into relevant minute blocks and time epochs. These analyses were integrated with the optogenetics manipulations and fiber photometry signals to establish the relationship between neural activity and behavior. We did not see differences in self administration behavior between the dLight and GCaMP recording groups, so behavioral data are aggregated in the main figures. Behavioral data disaggregated by sensor are shown in **Fig S2**. Further, we did not see consistent sex differences in either intermittent access to cocaine self-administration or BLA dopamine function, in line with our past studies (Collins et al., 2023; Brickner et al., 2026), and so for the main text figures male and females data are aggregated. Data disaggregated by sex are shown in **Fig S3**.

### Fiber photometry analysis

Photometry data was analyzed using custom MATLAB pipelines and GraphPad Prism similar to our previous studies (Engel et al., 2024; Stelzner et al., 2025; Brickner et al., 2026). First, the photometry signals (405 nm/415 nm isosbestic and 465 nm) were low pass filtered and downsampled to 40 Hz. The isosbestic signal was then aligned to the 465 nm signal by applying a least-squares linear fit and this fitted isosbestic signal was used to normalize the 465 nm signal. We calculated ΔF/F for dLight or GCaMP as follows: ΔF/F= (465 nm signal / fitted 415 nm signal)/(fitted 415 nm signal). Changes in ΔF/F fluorescence were Z-scored during time points of interest relative to a pre-event baseline.

Timestamps for each video recording (recorded at 10–20 FPS) were recorded in the Synapse photometry data file. Cue presentations, nose pokes, cocaine pump onset, and footshocks were assigned a TTL signal from Med-PC that was used to timestamp the photometry data file in the Synapse software. The normalized ΔF/F signals were extracted 5 s before and 10 s following each drug availability transition and Z-scored to the 5 s pre-transition period to account for drift in the signal across session duration. Normalized signals for nose pokes were extracted in the window from 2 s preceding to 10 s following each nose poke and Z-scored to the 2 s pre-nose poke period. Nose pokes were binned according to whether they occurred during the drug available or unavailable periods. Normalized signals for shock responses were extracted in the window from 2 s preceding to 10 s following each shock delivery and Z-scored to the 2 s pre-shock delivery. Photometry signal metrics, including the event-evoked peak Z-score and the area under the curve (AUC) were quantified and compared across training phases. AUC values were calculated from the Z-scored traces by numerical integration via the trapezoidal method using the trapz function (Matlab).

### Statistics

Behavior and photometry signal data were analyzed with a combination of ANOVA models (one-way, two-way, and mixed effects), t-tests, and Pearson correlations. Post-hoc comparisons and planned t-tests were used to clarify main effects and interactions, with a Bonferroni correction applied. Photometry traces represent average values of sites/subjects, not individual trials. The data are expressed as the mean ± s.e.m. All tests were twosided, and statistical significance was set at p<0.05.

### Histology

After experiments, rats were deeply anesthetized via intraperitoneal injections of Fatal-Plus (2ml/kg; Paterson Veterinary) before being transcardially perfused with cold phosphate buffered saline (PBS) and subsequent 4% paraformaldehyde (PFA). Brains were extracted and placed in 4% PFA solution for ∼24 hours, then transferred to a cryoprotectant solution of 30% sucrose in PBS for at least 48 hours. Tissue was sectioned at 50 μm on either a freezing microtome (ThermoFisher) or a cryostat (Leica CM1900). To validate proper viral expression as well as optic fiber placements, brain sections containing the BLA and VTA were mounted on microscope slides. The slides were cover-slipped with a Vectashield mounting medium containing the DAPI nuclear stain. Images of brain sections were taken on a fluorescent microscope (Keyence BZ-X710) with a 4x and a 10x air immersion objective. Photometry and/ or optogenetic data from each rat was only included in analysis if both the virus and fiber were located within the target region. Rats with mistargets of either metric were excluded. Virus spreads and fiber positions were compared to the Rat Brain Atlas (Paxinos and Watson, 2007) and visualized with Adobe Illustrator.

#### Immunohistochemistry

To visualize dopamine projections in the amygdala, we performed immunohistochemistry for tyrosine hydroxylase. Brain sections were washed in PBS and incubated with bovine serum albumin (BSA) and Triton X-100 (each 0.2%) for 20 min. 10% normal donkey serum (NDS) was added for a 30-min incubation, before primary antibody incubation (rabbit anti-TH, 1:500, Invitrogen) overnight at 4°C in PBS with BSA and Triton X-100 (each 0.2%). Sections were then washed and incubated with 2% NDS in PBS for 10 minutes and secondary antibodies were added (1:200 594 donkey anti-rabbit) for 2 hours at room temperature. Sections were washed 2 times in PBS and mounted with Vectashield containing DAPI before imaging.

## Supplemental Material

**Fig S1.**
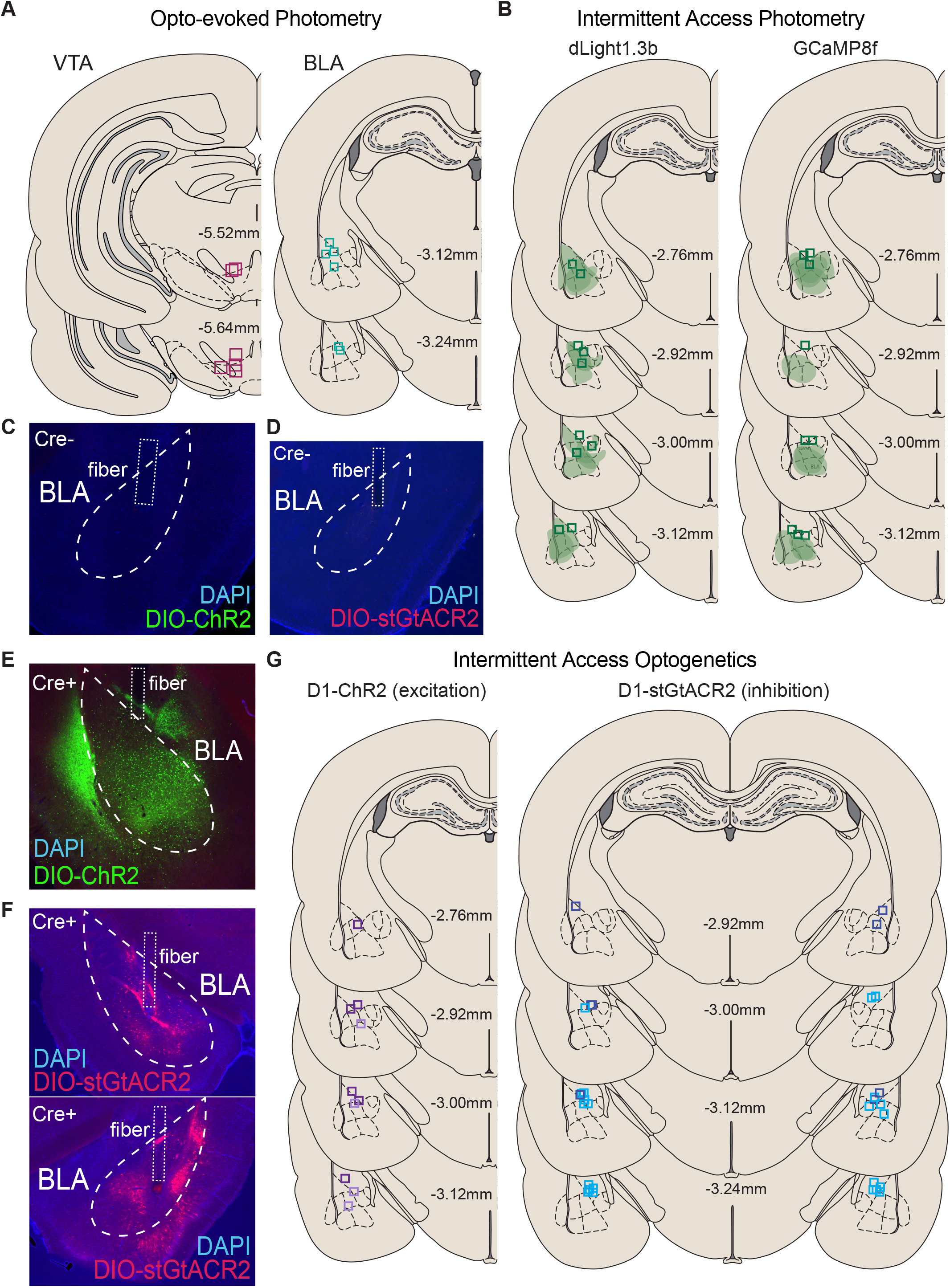
Histology for fiber photometry and optogenetics experiments. A) Optic fiber placements for rats (n=6) used to measure optogenetically-evoked dopamine signals in the BLA following optogenetic activation of VTA dopamine neurons. B) Viral spread and optic fiber placements for the photometry intermittent access experiments for the dLight1.3b (left, n=10) and GCaMP8f (right, n=9) rats. Representative images depicting lack of viral expression for the cre-negative control rats in the C) D1-ChR2 experiments and D) D1-stGtACR2 experiments. E) Representative histology images showing virus expression of the excitatory ChR2 opsin from a unilateral cre-dependent viral injection in the BLA of D1-cre rats. F) Representative histology images showing virus expression of the inhibitory stGtACR2 opsin from a bilateral cre-dependent viral injection in the BLA of D1-cre rats. G) Optic fiber placements for the optogenetic manipulation intermittent access experiments for the D1-ChR2 rats (left, n=10; active: light, control: dark) and the D1-stGtACR2 rats (right, n=14; active: light, control: dark).

**Fig S2.**
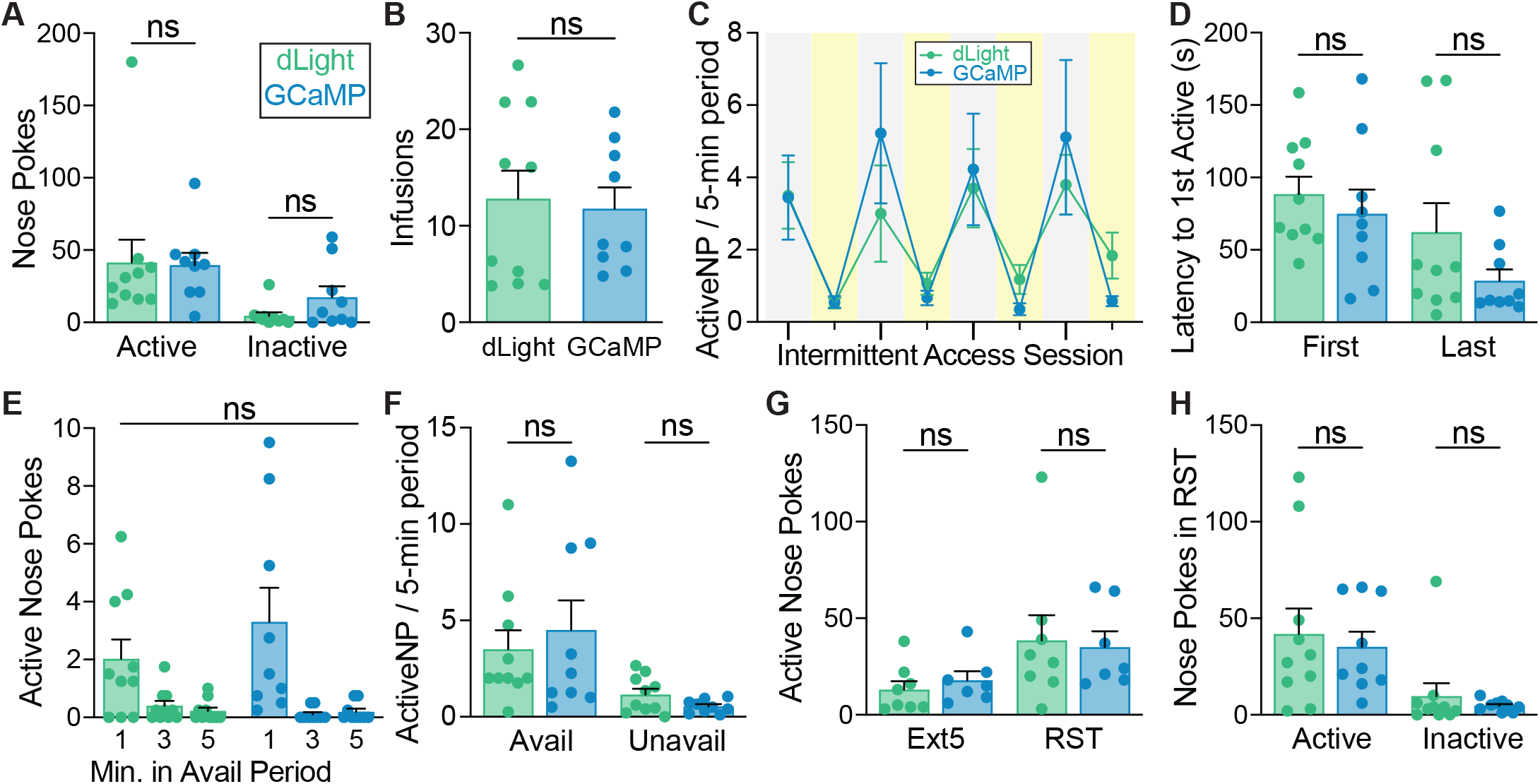
State-guided cocaine seeking behavior did not vary by photometry viral condition. A) Nose poke levels were not different between the dLight (green; n=10) and GCaMP groups (blue; n=9) during continuous access training day (F(1,17)=0.2861, p=0.5997). B) Rats in both viral groups self-administered cocaine at similar levels in terms of average daily cocaine infusions earned during intermittent access training (t(17)=0.2795, p=0.7832). C) dLight and GCaMP rats similarly discriminated between cocaine available and unavailable periods during intermittent access training and focused seeking bouts to available periods. Gray boxes represent drug available periods (lights off), while yellow boxes represent drug unavailable (lights on). D) The average latency to first active nose poke during the drug available periods was similar between virus conditions across intermittent access training (F(1,17)=3.361, p=0.0843). E) There were no significant main effects of virus on the temporal pattern of active nose pokes within the drug available periods during the final intermittent access session (F(1,17)=0.4507, p=0.5110). F) Average active and inactive nose pokes during the 5-min drug available periods and in the unavailable periods binned into 5-min blocks on the last intermittent access day split between the dLight and GCaMP viral conditions. Seeking behaviors in each period were similar (F(1,17)=0.03737, p=0.8490). G) Rats in both viral conditions who underwent extinction training (dLight: n=8; GCaMP: n=7) had similar active poke levels on extinction day 5 and the reinstatement test (F(1,13)=0.004804, p=0.9458). H) Rats (dLight: n=10; GCaMP: n=9) made a similar number of active and inactive nose pokes during cue-induced reinstatement, regardless of viral group (F(1,17)=0.3084, p=0.5859). Data represent mean ± SEM. ns: not significant. Avail: drug available. Unavail: drug unavailable. Ext: extinction. RST: reinstatement.

**Fig S3.**
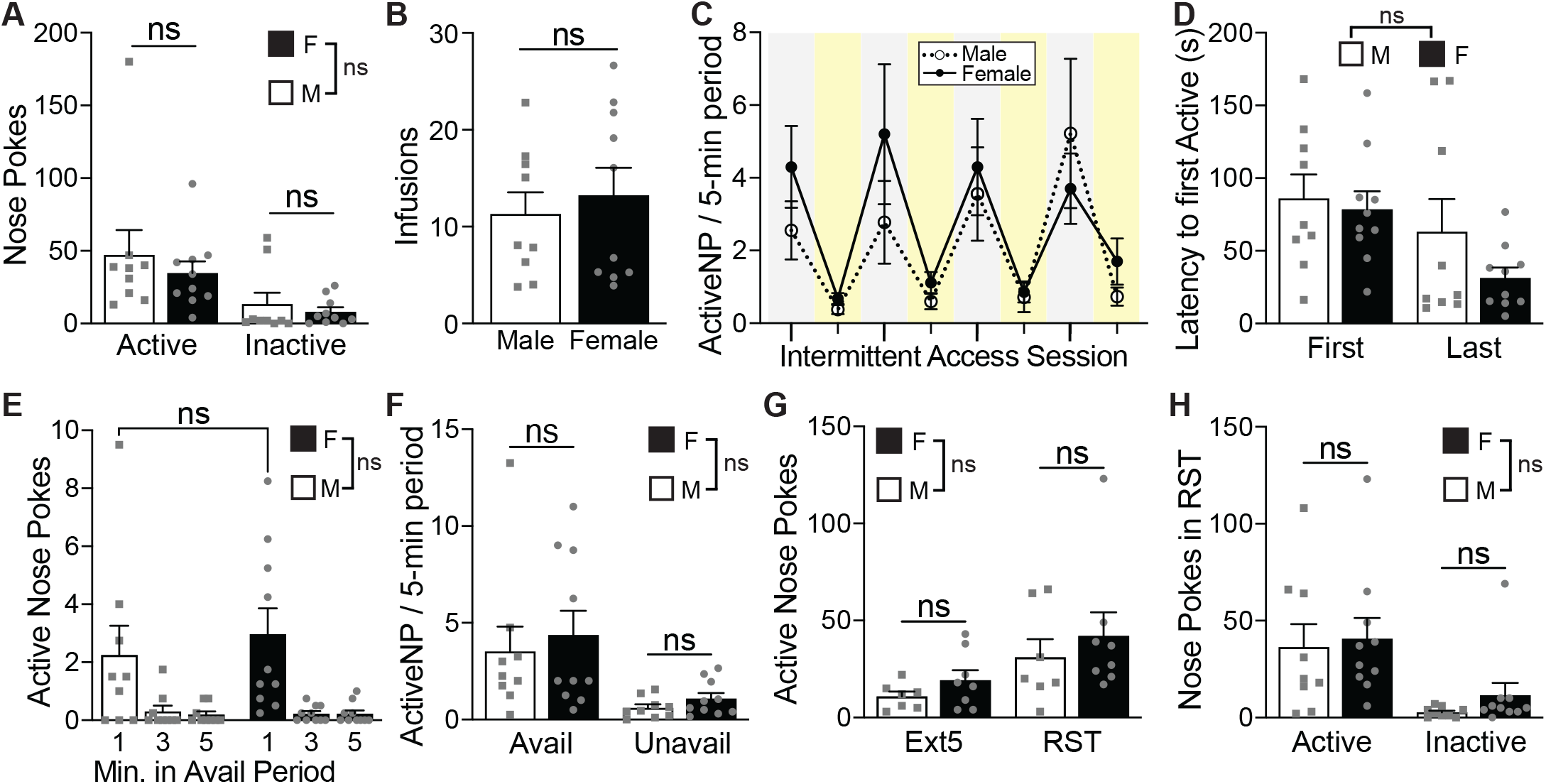
State-guided cocaine seeking behavior did not vary by sex. A) There was no difference in nose poke behavior between females (n=10; filled) and males (n=9; open) during the final continuous access training day (F(1,17)=0.7717, p=0.3919). B) Male and female rats self-administered cocaine at similar levels in terms of average daily cocaine infusions earned during intermittent access training (t(17)=0.5361, p=0.5988). C) Males and females focused their cocaine seeking bouts to the drug available periods to similar degrees across during intermittent access training. Gray boxes represent drug available periods (lights off), while yellow boxes represent drug unavailable (lights on). D) The average latency to first active poke during the drug available periods were comparable between sexes in the first and last intermittent access sessions (F(1,17)=2.162, p=0.1597). E) On the final intermittent access session, the temporal pattern of active nose pokes made within the drug available periods was not modulated by sex (F(1,17)=0.2194, p=0.6455), as females and males exhibited similar levels of active responding during minute one of the availability period (p=0.3628). F) Average active and inactive nose pokes during the 5-min drug available periods and in the unavailable periods binned into 5-min blocks on the last intermittent access training day split between the sexes. Seeking behaviors in each period were similar between males and females (F(1,17)=0.4488, p=0.5119). G) Female (n=8) and male (n=7) rats who underwent extinction training had similar numbers of total active pokes during the final extinction day and the reinstatement test (F(1,13)=1.266, p=0.2808). H) Nose poke levels did not differ between females (n=10) and males (n=9) during cue-induced reinstatement (F(1,17)=0.3797, p=0.5460). Data represent mean ± SEM. *p<0.05, **p<0.01, ns: not significant. Avail: drug available. Unavail: drug unavailable. Ext: extinction. RST: reinstatement.

**Fig S4.**
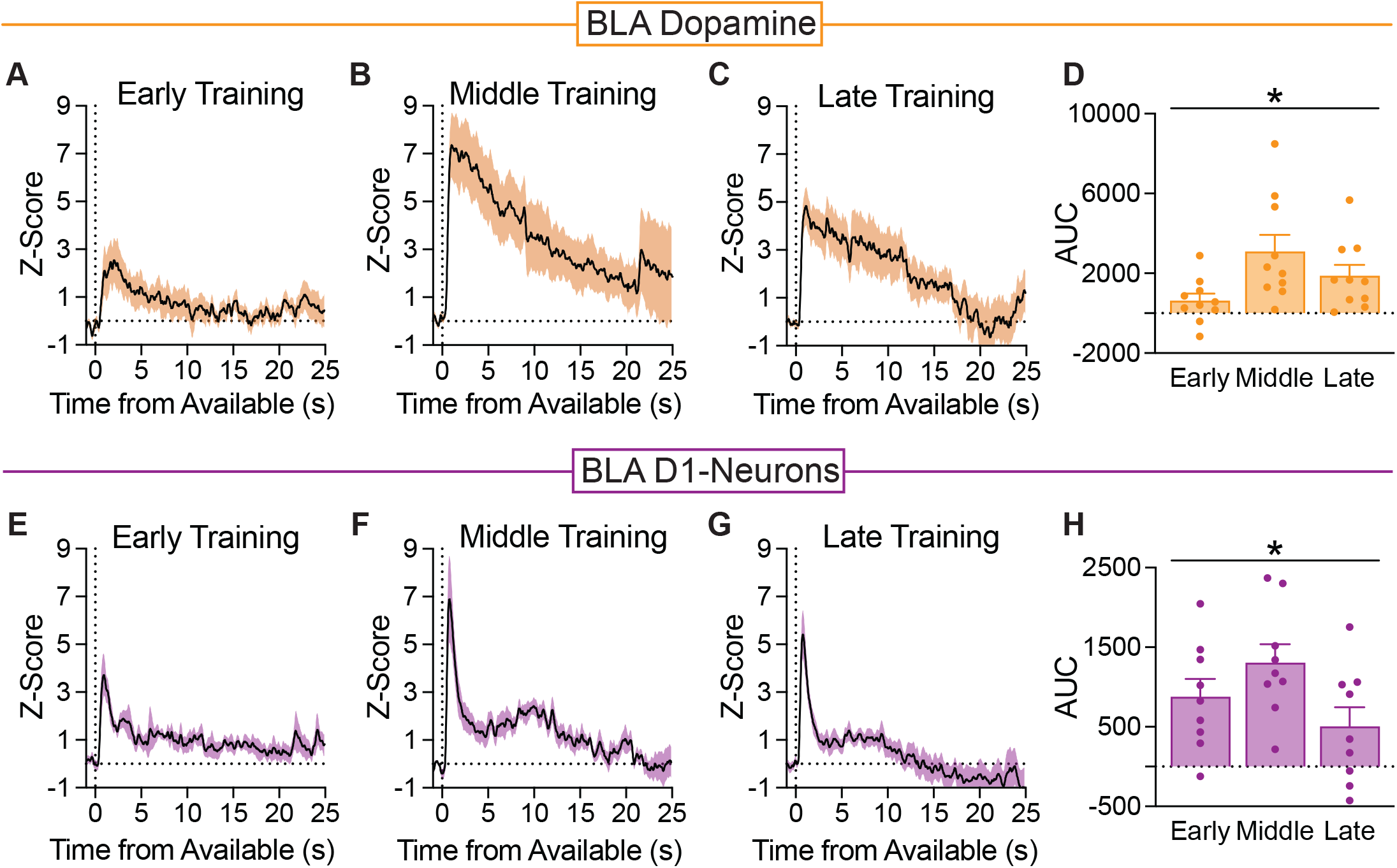
Drug availability state transitions evoke unique temporal patterns of BLA dopamine signaling versus D1 neuron activity. A-C) The BLA dopamine response to the first 25s of the drug availability state onset in rats (n=10) during intermittent access training. The availability state dopamine signal became more sustained across the A) early, B) middle, and C) late phases of intermittent access. D) These dopamine signals dynamically changed in magnitude across training (repeated measure, one-way ANOVA, F(2,18)=4.912, p=0.0198). E-G) The BLA D1-neuronal response to the first 25s of the drug availability state onset in rats (n=9) across intermittent access training. The availability state evoked rapid, phasic D1-neuron activity across E) early, F) middle, and G) late intermittent access training. H) These D1-neuron signals dynamically changed in magnitude across training (repeated measure, one-way ANOVA, F(2,16)=3.817, p=0.0441). Data represent mean ± SEM. *p<0.05. AUC: Area under curve.

**Fig S5.**
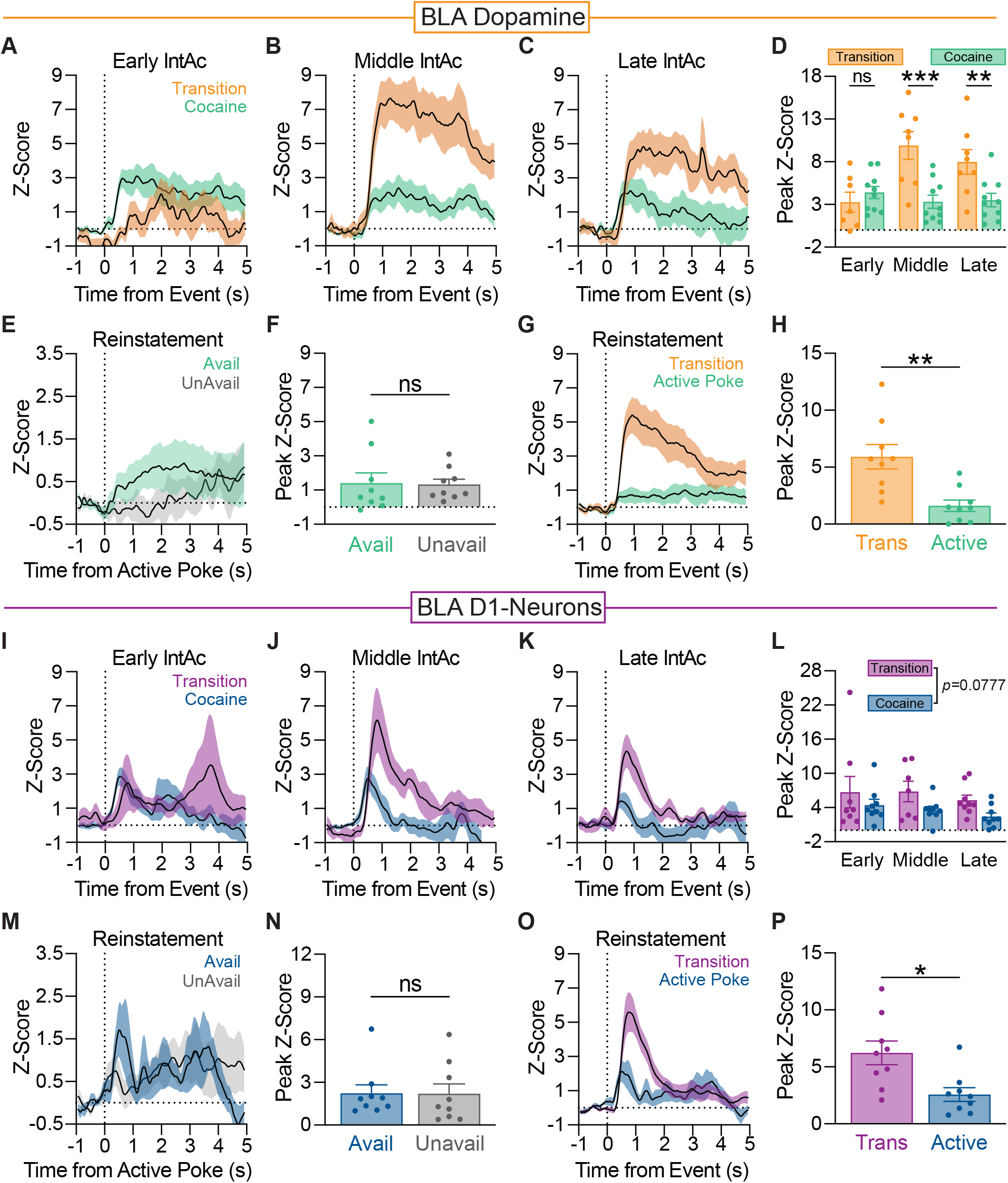
Dopamine and D1-neuron responses to drug availability state transitions before cocaine is on board. A) Comparisons between BLA dopamine signals (n=10) evoked by the first drug availability state transition and the first cocaine-paired active nose poke in early, B) middle, and C) late intermittent access training. D) In general, the first drug available state shift prompted larger dopamine events compared to the first cocaine-seeking action in intermittent access (mixed-effects ANOVA, main effect of event: F(1,9)=16.53, p=0.0028), with the different event types prompting different dopamine levels as intermittent access training progressed (training x event interaction: F(2,11)=7.483, p=0.0089) during middle (p=0.0001) and late (p=0.0044) training, where the availability transition evoked larger dopamine signal. E) Small BLA dopamine signals were associated with active nose pokes made in the “drug available” and “drug unavailable” periods in the reinstatement test, and F) these signals were not different (t(8)=0.9194, p=0.9194). G) In contrast, large BLA dopamine signals were evoked by the reintroduction of the “drug available” transitions in the reinstatement test, which were larger than signals associated with active nose pokes (t(8)=3.985, p=0.0040). I) Comparisons between BLA D1-neuron responses (n=9) to the first drug availability state transition and the first cocaine-paired active nose poke in early, J) middle, and K) late intermittent access training. L) The peak signal for the availability state transition trended larger than the cocaine-taking signal throughout intermittent access training (mixed-effects ANOVA, main effect of event: F(1,8)=4.093, p=0.0777). M) BLA D1-neurons were modestly active after rats (n=9) made active nose pokes during both the “drug available” and “unavailable” periods in cue-maintained reinstatement, and N) these signals were not different (t(8)=0.07123, p=0.9450). O) In contrast, larger BLA D1-neurons signals were evoked by the reintroduction of the “drug available” transitions in the reinstatement test, P) relative to signals associated with active nose pokes (t(8)=2.937, p=0.0188). Data represent mean ± SEM. *p<0.05, **p<0.01, ***p<0.001, ****p<0.0001, ns: not significant.

## Notes

### Competing Interest Statement

The authors have declared no competing interest.

